# NFIC regulates ribosomal biology and ER stress in pancreatic acinar cells and suppresses PDAC initiation

**DOI:** 10.1101/2021.08.09.455477

**Authors:** Isidoro Cobo, Sumit Paliwal, Júlia Melià-Alomà, Ariadna Torres, Jaime Martínez-Villarreal, Fernando García, Irene Millán, Natalia del Pozo, Joo-Cheol Park, Ray J. MacDonald, Javier Muñoz, Francisco X. Real

**Affiliations:** Epithelial Carcinogenesis Group, Spanish National Cancer Research Centre-CNIO, Madrid, Spain; CIBERONC, Madrid, Spain; Departament de Ciències Experimentals i de la Salut, Universitat Pompeu Fabra, Barcelona, Spain; Proteomics Unit, Spanish National Cancer Research Centre-CNIO, Madrid, Spain. ProteoRed - ISCIII; Department of Oral Histology-Developmental Biology, School of Dentistry, Seoul National University, Seoul, Korea; Department of Molecular Biology, University of Texas Southwestern Medical Center, Dallas, TX, USA

**Keywords:** NFIC, pancreas, acinar differentiation, ribosome, endoplasmic reticulum stress, unfolded protein response, transcriptional networks, pancreatitis, pancreatic cancer

## Abstract

Tissue-specific differentiation is driven by specialized transcriptional networks. Pancreatic acinar cells crucially rely on the PTF1 complex, and on additional transcription factors, to deploy their transcriptional program. Here, we identify NFIC as a novel regulator of acinar differentiation using a variety of methodological strategies. NFIC binding sites are found at very short distances from NR5A2-bound genomic regions and both proteins co-occur in the same complex. *Nfic* knockout mice show reduced expression of acinar genes and, in ChIP-seq experiments, NFIC binds the promoters of acinar genes. In addition, NFIC binds to the promoter of, and regulates, genes involved in RNA and protein metabolism; in *Nfic* knockout mice, p-RS6K1 and p-IEF4E are down-regulated indicating reduced activity of the mTOR pathway. In 266-6 acinar cells, NFIC dampens the ER stress program through its binding to ER stress gene promoters and is required for complete resolution of Tunicamycin-mediated ER stress. Normal human pancreata from subjects with low NFIC mRNA levels display reduced epxression of genes down-regulated in *Nfic* knockout mice. Consistently, NFIC displays reduced expression upon induced acute pancreatitis and is required for proper recovery after damage. Finally, expression of NFIC is lower in samples of mouse and human pancreatic ductal adenocarcinoma and *Nfic* knockout mice develop an increased number of mutant *Kras*-driven pre-neoplastic lesions.

## INTRODUCTION

Pancreatic acinar cells are highly specialized protein synthesis factories that have a well-developed rough endoplasmic reticulum (ER), a prominent Golgi complex, and abundant secretory granules^1^. Acinar differentiation is contingent on the activity of a master regulator, the adult PTF1 complex, composed of the pancreas-specific transcription factors (TFs) PTF1A and RPBJL and the ubiquitous protein E47^2,3^. PTF1 binds the proximal promoter of genes coding for digestive enzymes, secretory proteins and other TFs, and activates their expression. The PTF1 complex is the main driver of acinar differentiation but additional TF with tissue-restricted expression patterns are implicated in the fine-tuning of this process, including GATA6^4^, MIST1^5^, and NR5A2/LRH-1^6,7^. Acinar cells play a crucial role in acute and chronic pancreatitis, two common and disabling conditions. Recent work using genetic mouse models has shown that, upon expression of mutant *KRas*, acinar cells can be the precursors of Pancreatic Intraepithelial Neoplasia (PanIN) and pancreatic ductal adenocarcinoma (PDAC)^8,9^.

Our laboratory and others have shown that the acinar differentiation program acts as a tumor suppressor in the pancreas. Monoallelic or homozygous inactivation of several acinar transcriptional regulators in the germline, the embryonic pancreas, or the adult pancreas can result in compromised acinar function that favors loss of cellular identity and poises acinar cells for transformation upon activation of mutant KRas^10,11,12^. The tumor suppressive function of these TF is not obvious because the exocrine pancreas has a large functional reserve, i.e. massive alterations in cellular function need to occur in order to be reflected in histological or clinical changes.

Here, we use bioinformatics tools to identify NFIC as a novel acinar regulator. NFIC is a member of the nuclear factor I family of TFs that regulate both ubiquitous and tissue-restricted genes^13^. In the mammary gland, NFIC activates the expression of milk genes involved in lactation^14^. Furthermore, it acts as a breast cancer tumor suppressor, as it directly represses the expression of *Ccnd1* and *Foxf1*, a potent inducer of epithelial-mesenchymal transition (EMT), invasiveness, and tumorigenicity. Additional roles have been proposed through the regulation of *Trp53*^15,16,17^. The physiological role of NFIC has been best studied in dentinogenesis, since *Nfic*^-/-^ mice develop short molar roots and display aberrant odontoblast differentiation and dentin formation^18^. NFIC regulates odontoblast-related genes, including *Dssp*^19^, Wnt^20^, and hedgehog signaling^21^.

Using a combination of omics analyses and studies in knockout mice and cultured cells, we now uncover novel roles of NFIC as a regulator of acinar function whose major impact is at the level of the ER stress response in murine and human pancreas. Unlike most other TFs previously identified as required for full acinar function, NFIC belongs to a novel family of acinar regulators with tissue-wide expression. NFIC dysregulation sensitizes the pancreas to damage and neoplastic transformation.

## RESULTS

### Identification of novel transcription factors involved in the regulation of pancreatic acinar differentiation

To discover novel transcription factors that might cooperate with known acinar regulators (e.g. PTF1A, GATA6, NR5A2, and MIST1), we reanalyzed publicly available ChIP-sequencing data and used HOMER to search for motifs enriched in the sequencing reads. As expected, the cognate binding sites of these factors were the top enriched motif in each respective analysis (Figure 1A). Motifs corresponding to RBPJ/RBPJL and HNF1, known regulators of acinar differentiation, were also enriched, thus validating the strategy applied. In addition, we found consistent enrichment of the NF1(CTF)/NFIC motif across all the experiments, with lowest p-values in the NR5A2 ChIP-Seq dataset. This motif is significantly enriched in NR5A2 ChIP-Seq peaks from normal adult mouse pancreas^6^ (Figure 1A) but not in those from E17.5 pancreas^7^ nor from mouse ES cells^22^, pointing to temporal and lineage identity specificity (Figure 1B). Of all NFI family members, NFIC is expressed at highest levels in both mouse and human pancreas (Figure 1C); therefore, we focused on NFIC for further study. Using immunoprecipitation and western blotting, we found that NFIC and NR5A2 are present in the same complex in normal adult pancreas - but not in E17.5 pancreas (Figure 1D). Analysis of the spacing between NR5A2 and NFIC motifs in the genomic regions bound by NR5A2, using the SpaMo tool from MEME suite, showed that NFIC binding motifs are located in close proximity to NR5A2 binding motifs, with a spacing of 29 nucleotides being the most significantly conserved distance (P=e-16) (Figure 1E). Analysis of the published ChIP-Seq data revealed several PTF1A and NR5A2 peaks in the proximal *Nfic* promoter and the binding was confirmed by ChIP-qPCR (Figure 1F), strongly suggesting that *Nfic* is a PTF1A and NR5A2 target. Using ChIP-qPCR, NFIC was found to bind the promoter of *bona fide* acinar genes such as *Cela2a, Cpa1, Ctrb1, Pnlip* and *Nrob2* that were similarly bound by NR5A2 (Figure 1G). The above observations support the notion that NFIC is a novel pancreatic acinar transcription factor network member.

**Figure 1.**
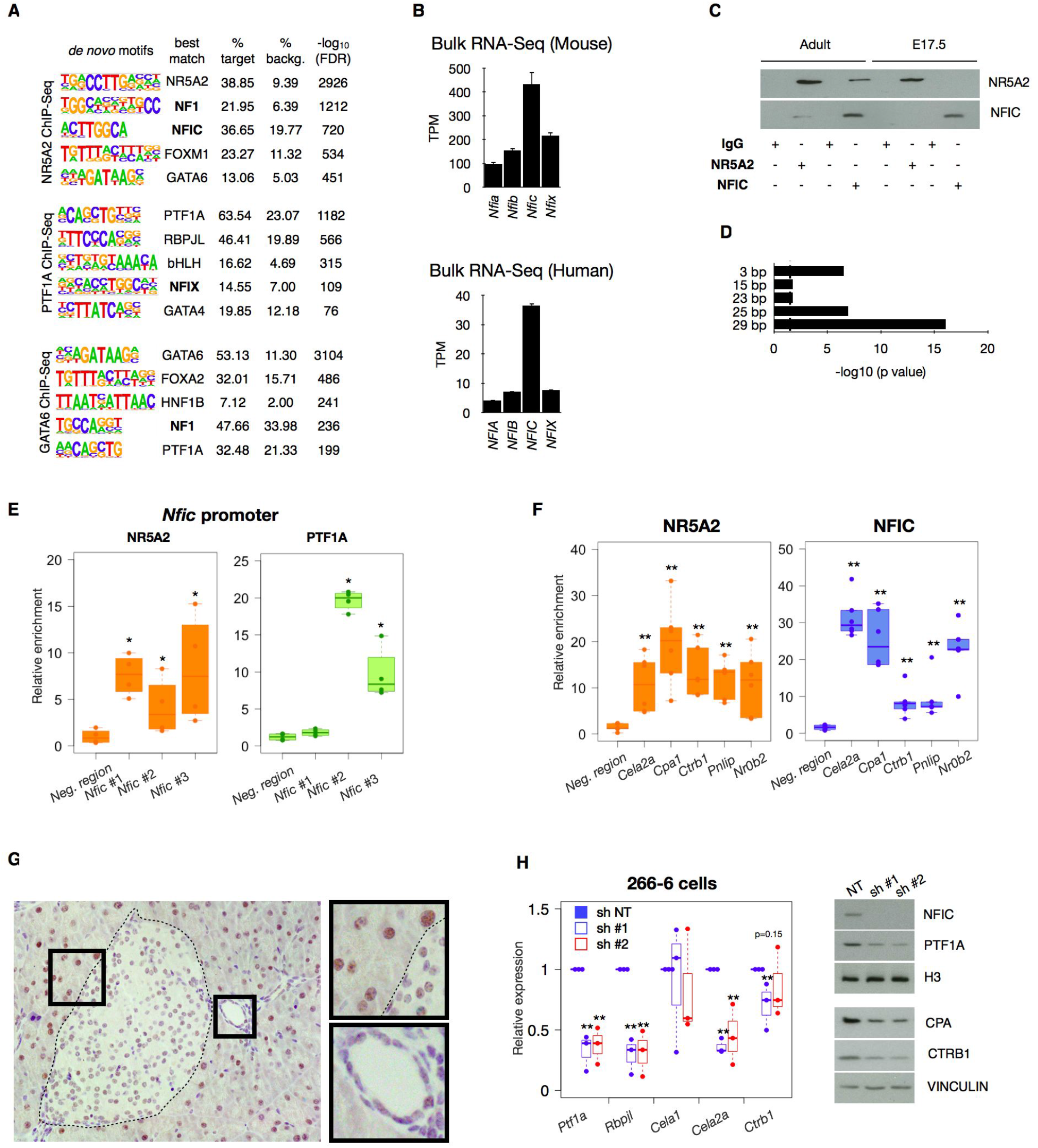
The NR5A2 is a novel acinar regulator. (A) HOMER *de novo* motif analysis for NR5A2, PTF1A and GATA6 ChIP-Seq in mouse pancreata showing enrichment in NF1/NFI motifs. (B) Expression of *NFI* transcripts in mouse (upper panel) or human (lower panel) pancreata assessed by RNA-Seq showing that *NFIC* is the family member expressed at highest levels. (C) Immunoprecipitation-western blotting analysis showing that NFIC and NR5A2 are part of the same complex in adult, but not in embryonic, pancreas. (D) Spamo analysis showing distance conservation of the NR5A2 and NFIC motifs in the regions bound by NR5A2. (E) ChIP-qPCR of NR5A2 and PTF1A binding at the *Nfic* promoter (one region in NR5A2 peak1 and two regions in NR5A2 peak3); controls as in panel F (n=4/group). (F) ChIP-qPCR of NR5A2 and NFIC binding to the promoter of digestive enzyme genes and *Nr0b2*, compared to a control (Neg) region (normalized to unrelated IgG) (n=6/group). (G) IHC analysis of NFIC in normal adult mouse pancreas showing higher expression in acinar cells and lower expression in endocrine and ductal cells (insets). (H) Lentiviral *Nfic* knockdown in 266-6 cells showing reduced expression of transcripts coding for digestive enzyme transcripts and pancreatic TFs (RT-qPCR) (left panel); western blotting analysis of the corresponding samples interfered with non-targeting (NT) or *Nfic*-targeting shRNAs (n=3).

To assess the cellular distribution of NFIC, we performed immunohistochemistry (IHC) with a well-validated antibody. In normal 8 week-old pancreas, NFIC is expressed at high levels in acinar cells and at lower levels in endocrine and ductal cells (Figure 1H). These results were validated using triple immunofluorescence (IF) with antibodies detecting PTF1A, INS1, and KRT19 (Supplementary Figure 1). In contrast, NFIC was undetectable at E12.5 and E14.5 in CDH1^+^;PTF1A^+^ pancreatic progenitors (Supplementary Figure 2A,B) but it was detected at E16.5 and E18.5 in PTF1A^+^ acinar as well as in KRT19- and INS1-expressing cells (Supplementary Figure 2C,D).

**Figure 2.**
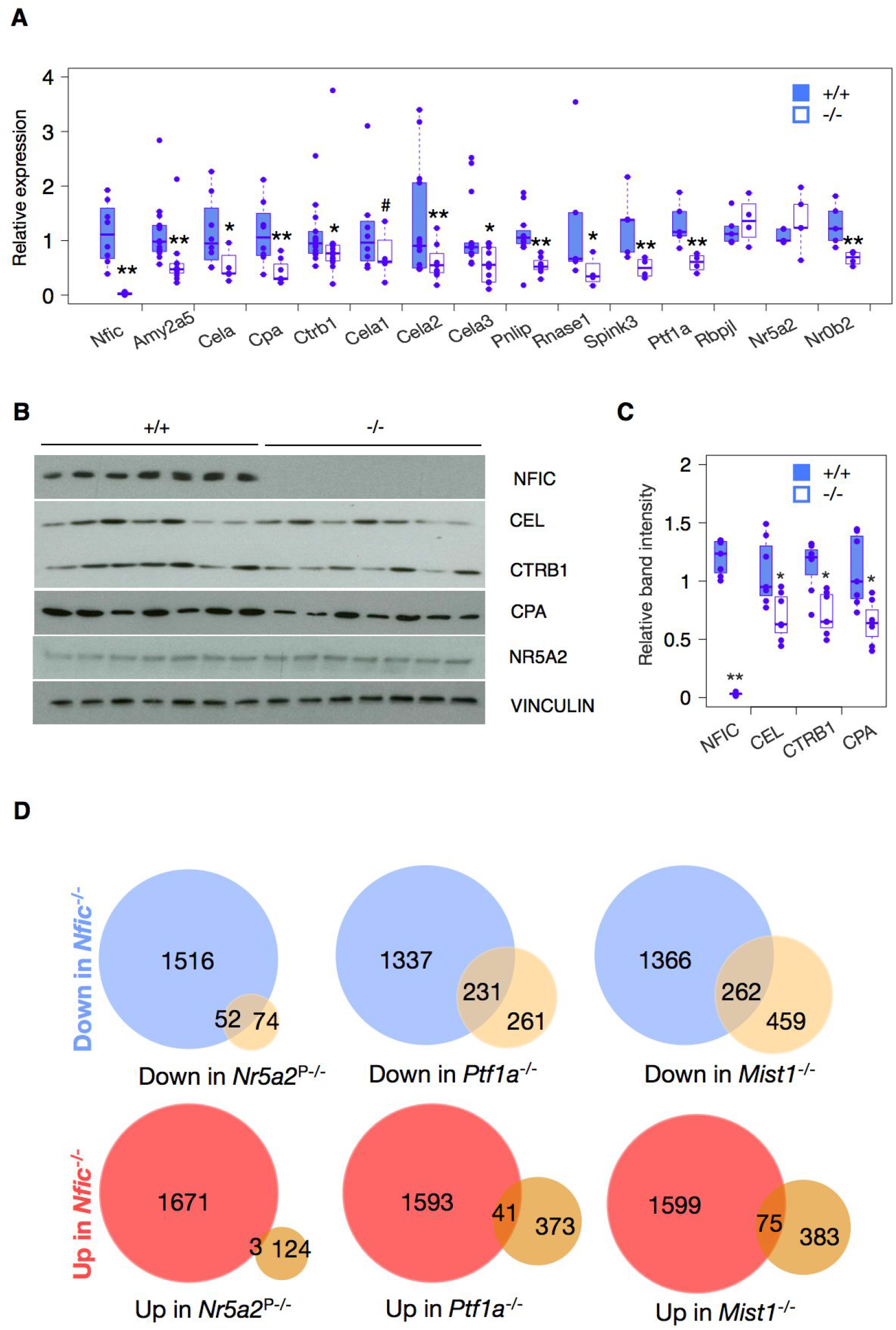
NFIC is required for normal acinar cell differentiation. (A) RT-qPCR showing reduced expression of transcripts coding for digestive enzymes and pancreatic TF in *Nfic*-/-pancreata. (B) Western blotting showing reduced expression of digestive enzymes in *Nfic*^-/-^ pancreata (n=7/group). (C) Densitometric quantification of panel 3B: band intensity normalised to loading control, relative to wild-type pancreata. (D) Comparison of the overlap of DEG in the pancreas of *Nfic*^-/-^ vs. that of mice lacking NR5A2, PTF1A, and MIST1 (details in text). Statistics: two-tailed Student-T test. Significant overlap is shown for down-regulated genes compared to a random list of genes. “N-1” chi-squared test was used to calculate statistical significance.

To determine whether NFIC is required for the expression of digestive enzyme transcripts, we first knocked down *Nfic* in 266-6 acinar cells using lentiviral shRNAs: a significant down-regulation of *Ctrb1* and *Cela2a* - as well as *Ptf1a* and *Rbpjl* - was demonstrated. Expression of PTF1A, CTRB1, and CPA proteins was similarly reduced (Figure 1H), suggesting an important role for NFIC in the regulation of late stages of acinar differentiation.

### NFIC is part of the transcriptional network responsible for acinar identity and function

To further assess the role of *Nfic* in pancreatic development and homeostasis, we used constitutive *Nfic* knockout mice^18^. *Nfic*^-/-^ mice are viable and have a normal weight at 8 weeks (not shown. *Nfic*^-/-^ pancreata appeared histologically normal and we did not find major differences in the expression of INS1, KRT19, and SOX9 (Supplementary Figure 3A), indicating that NFIC is not crucially required for pancreas development or differentiation. Results of glucose tolerance tests were similar in 8 week-old control and *Nfic*^-/-^ mice, except that glucose levels were reduced by 60-120 min in the latter (Supplementary Figure 3B,C).

**Figure 3.**
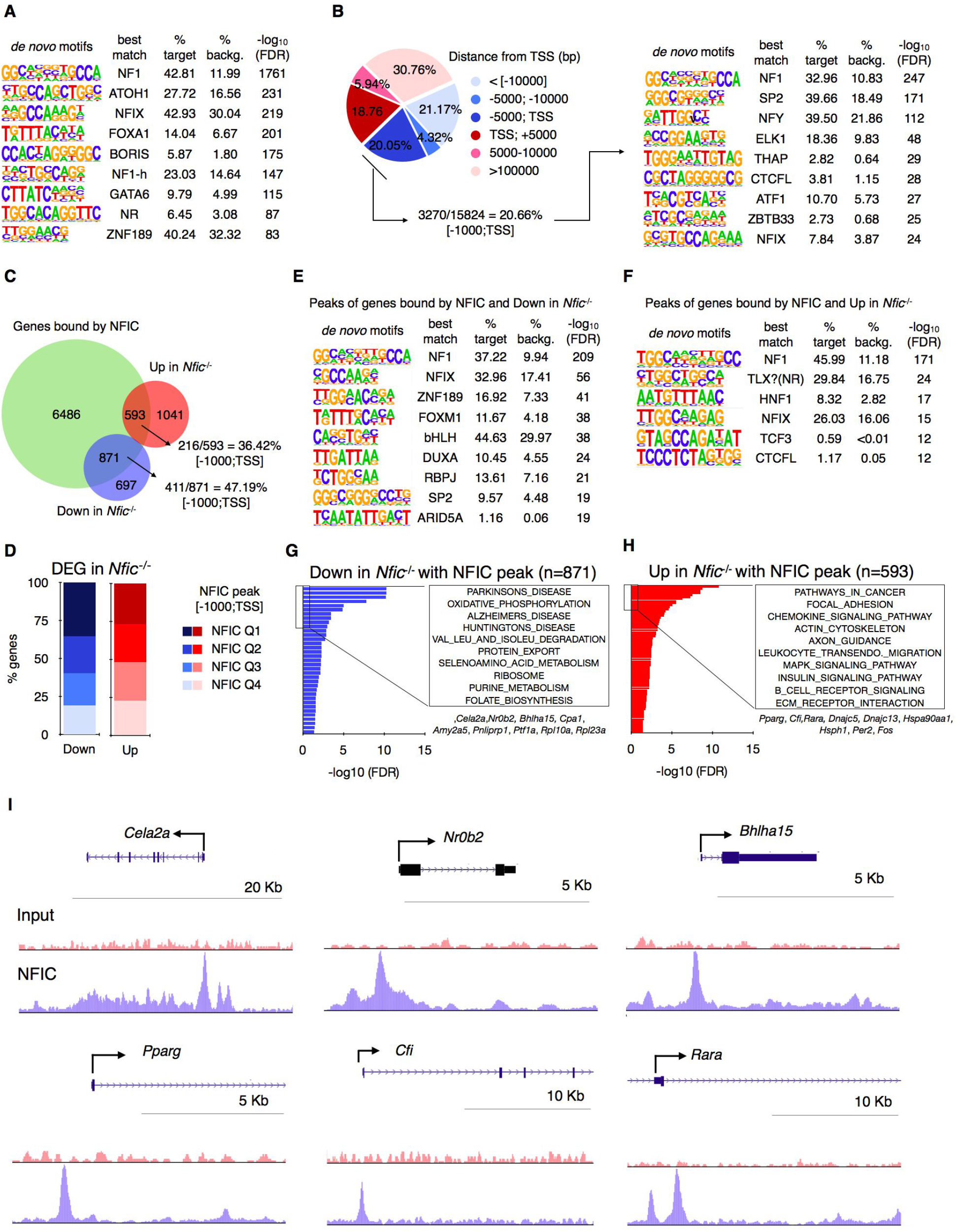
NFIC binds to genomic regions associated to genes involved in acinar differentiation, ER stress, UPR, and inflammation. (A) *De novo* motif analysis of NFIC ChIP-Seq showing NFI as the top-motif. (B) Distribution of NFIC ChIP-Seq peaks showing enrichment in regions close to the TSS (left) and the corresponding enrichment of the NFI, ELK and CTCF motifs (right). (C) Venn diagram showing the overlap between genes with an NFIC peak and those de-regulated in the *Nfic-/-* pancreas showing a greater overlap for the down-regulated genes. (D) Bar graph of the distribution of NFIC ChIP-Seq peaks based on score intensity and the overlap with genes de-regulated in *Nfic*-/-pancreata showing slight greater overlap in Q1, Q2 for the down-regulated genes. (E,F) Motif analysis of genes with an NFIC peak that are down-regulated (E) or up-regulated (F) in *Nfic*-/-pancreata; NFI is the top motif in both groups. (G,H) Gene set enrichment analysis of genes bound by NFIC and down-regulated (G) or up-regulated (H) in *Nfic*-/-pancreata showing down-regulation of *bona fide* acinar, ribosomal, and metabolic genes; and up-regulation of inflammatory, UPR, and ER stress genes. Boxes show representative examples of genes included in each analysis. (I) UCSC browser shots of NFIC ChIP-Seq showing enrichment for *Cela2a, Nr0b2, Bhlha15, Pparg, Cfi* and *Rara*.

Because histology lacks sensitivity to disclose subtle alterations in exocrine function^4,10,23^ we performed RNA-Seq of pancreata from 8-10 week-old wild type and *Nfic*^-/-^ mice. We identified 1641 and 1568 transcripts that were significantly up- and down-regulated, respectively, in *Nfic*^-/-^ pancreata. Multiple genes belonging to the exocrine differentiation program were among the down-regulated transcripts (e.g. *Ptf1a*, several digestive enzymes, and the acinar-specific kinase *Mknk1*^24^ and the differences were confirmed using qRT-PCR (Figure 2A) and at the protein level (Figure 2B,C, Supplementary Figure 4A,B). The RNA-Seq data also showed reduced expression of genes involved in epithelial polarity (e.g. *Muc1*) and cell adhesion (e.g. *Cdh1*) and up-regulation of transcripts coding for EMT markers (e.g. Vimentin, Twist, and N-cadherin/*Cdh2*; not shown). The down-regulation of *Cdh1* is in accordance with findings in dentinogenesis^19^. NR5A2 expression was similar in control and *Nfic*^*-/-*^ pancreata (Figure 2B), indicating that the effects of *Nfic* inactivation are not secondary to changes in NR5A2 expression.

**Figure 4.**
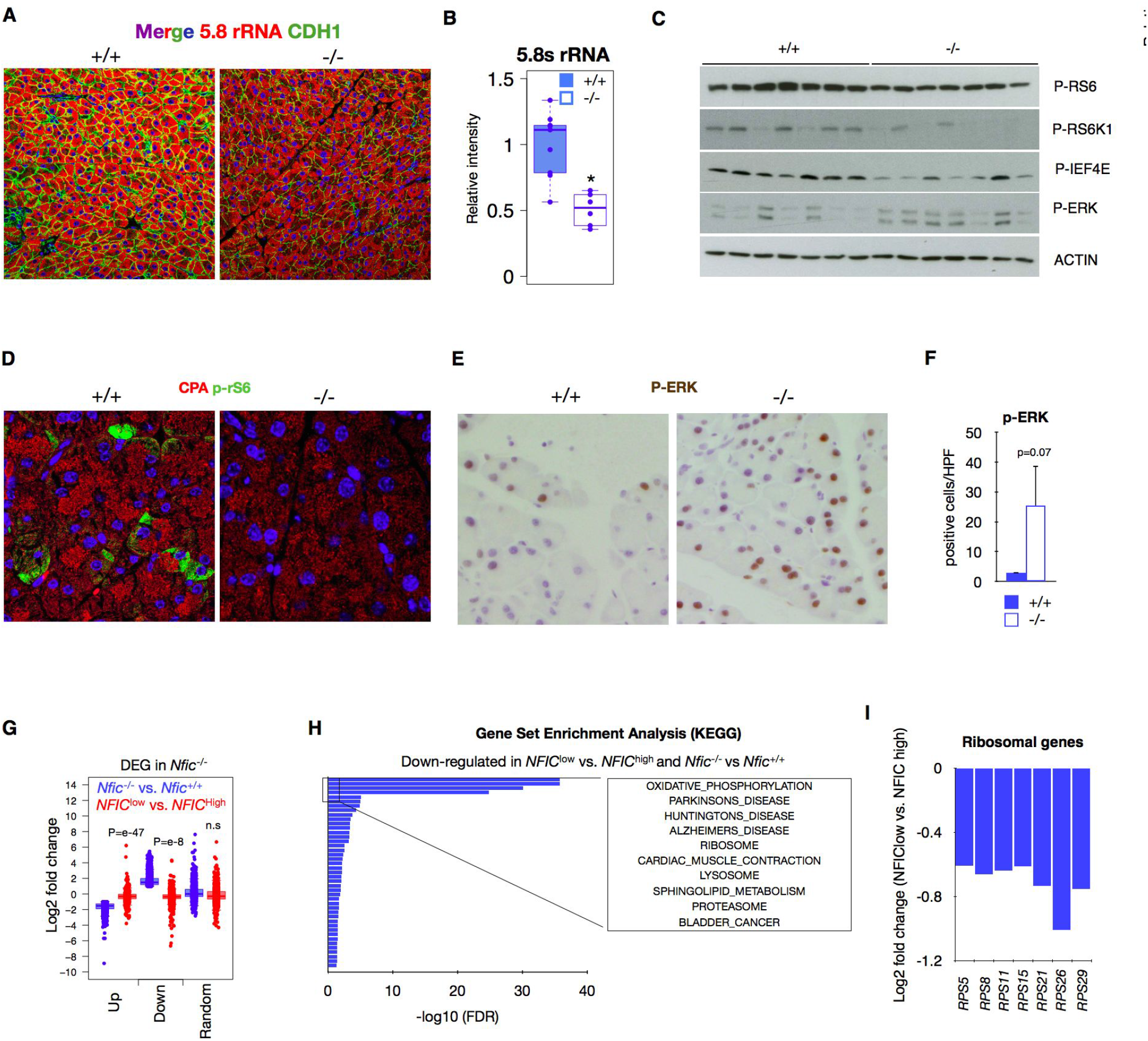
NFIC regulates protein biosynthesis in the pancreas. (A) IF analysis of reactivity with an antibody recognizing 5.8S rRNA and CDH1 shows decreased expression of both in *Nfic*^-/-^ acinar cells. (B) Quantification of panel 4A. (C) Western blotting showing mTOR and ERK pathway signaling changes in *Nfic*^-/-^ pancreata. (D) IF displaying down-regulation of CPA and P-RS6 in *Nfic*^-/-^ pancreata (n=2). (E,F) IHC analysis (E) of P-ERK and quantification (F) showing increased number of positive acinar cells in *Nfic*^-/-^ pancreata (n=2). (G) Boxplot plot showing the relationship between the expression of up-regulated, down-regulated, or a random set of genes, in control *Nfic*^*+/+*^ vs. *Nfic*^*-/-*^ mice and in histologically normal human pancreatic tissues samples (top 10 low-vs. top 10 high-expressing *NFIC* mRNA levels, as determined by RNA-Seq analysis [*NFIC*^low^ vs. *NFIC*^high^]). Data shows the concordant pattern between down-regulated genes in *Nfic*^-/-^ mice and *NFIC*^low^ human pancreata. “N-1” chi-squared test was used to calculate statistical significance. P-value was calculated comparing to a random gene list. (H) GSEA for genes that are concurrently down-regulated in *Nfic*^-/-^ vs. wild type pancreata and in *NFIC*^low^ vs. *NFIC*^high^ human pancreata. Genes were computed with KEGG data sets showing the similarities with those gene sets under-represented in *Nfic*^-/-^ mice (G). (I) Bar plot displaying the down-regulation of ribosomal genes in *NFIC*^low^ vs. *NFIC*^high^ human pancreata (P<0.001).

Up-regulated transcripts were enriched in inflammatory/immune gene sets, including chemokines (e.g. *S100a8, S100a9, Ccl5, Ccl7, Cxcl12, Cxcl3*) and complement components (e.g. *C1qb, C3, Cfb, Cfd*) (Supplementary Figure 5A). Selected changes were validated by RT-qPCR in total pancreas and in freshly isolated acini (Supplementary Figure 5B). These genes have putative *NFI* binding sites in their promoter region [-950bp; +50bp] (Supplementary Figure 5C). The up-regulation of inflammatory gene transcripts was accompanied by a 2-fold increase of CD45^+^ cells and Ki67^+^ acinar cells (Supplementary Figure 5D-F). These results suggest that NFIC contributes to restrain an inflammatory program in the pancreas.

**Figure 5.**
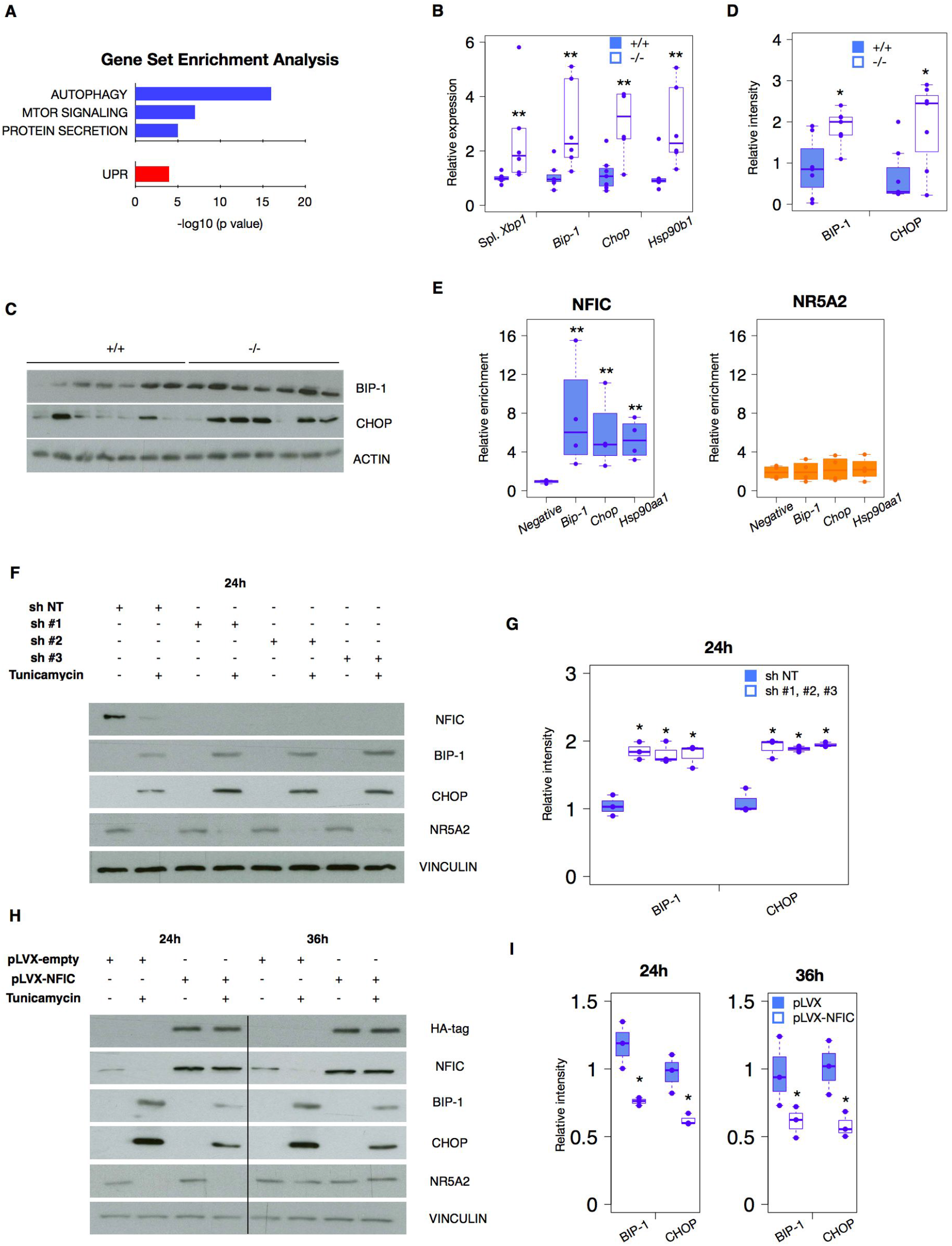
NFIC regulates aspects of UPR and ER stress resolution. (A) GSEA analysis of the UPR and ER stress gene sets^16^ down-regulated in *Nfic*^-/-^ pancreata and up-regulation of UPR. (B) RT-qPCR showing expression of spliced *Xbp1, Chop* (*Ddit3*), *Bip-1(Hspa5)*, and *Hsp90b1* in wild-type and *Nfic*^-/-^ pancreata (n>4/group). (C) Western blotting showing up-regulation of BIP-1 and CHOP in *Nfic*^-/-^ pancreata (n=7/group). (D) Densitometric quantification of data in panel 5C. (E) ChIP-qPCR showing binding of NFIC, but not NR5A2, to the promoters of *Hspa5*/*Bip*-1, *Ddit3* and *Hsp90aa1*. (F) Up-regulation of BIP-1 and CHOP in 266-6 cells treated with TM upon *Nfic* knock-down. (G) Quantification of data shown in 5F. (H) Reduced BIP-1 and CHOP expression in control and NFIC-overexpressing 266-6 cells treated with TM. (I) Quantification of data shown in 5H.

We quantified the overlap of differentially expressed genes in *Nfic*^*-/-*^ pancreata with that in mice in which *Nr5a2*^6^, *Ptf1a*^25^ or *Mist1*^26^ has been inactivated in the pancreas (PKO). We found a significant overlap of genes down-regulated in the pancreas of *Nfic*^*-/-*^ and *Nr5a2* pancreas-knockout, *Ptf1a* pancreas-knockout, or *Mist1* knockout mice [41% (52/126), 46.97% (231/492), 57.08% (262/459), respectively] but not of the up-regulated genes [2.36% (3/127), 9.99% (41/414), 16.37% (75/458), respectively] (Figure 2D) These findings strongly suggest that NFIC is a novel member of the acinar transcription factor network.

To identify direct NFIC target genes we performed ChIP-seq using pancreata from 8-10 week-old wild type mice. A total of 15824 peaks bound by NFIC were identified, corresponding to 9086 genes, with enrichment of motifs corresponding to NF1, ATOH1, and TF involved in acinar cell differentiation such as FOXA1, GATA6, and nuclear receptors, among others (Figure 3A). NFIC peaks were enriched in the vicinity of the TSS of genes [-1000; TSS, 22.66%] and these peaks displayed significant enrichment of NF1, SP2, THAP, ELK1 and AP-1 motifs (Figure 3B).

RNA-seq and ChIP-seq data were integrated to unveil genes/pathways directly regulated by NFIC: 36.3% (593/1634) of the differentially up-regulated genes (P< 0.05) and 55.54% (871/1568) of the down-regulated genes in *Nfic*-/-pancreata were bound by NFIC (P < 0.05). A greater percentage of down-regulated genes (47.19%, 411/871) relative to up-regulated genes (36.42%, 216/593), had NFIC peaks in the putative promoter region [-1000:TSS] (Figure 3C). The proportion of NFIC high affinity peaks as defined by the top two quartiles of peak score (Q1+Q2) was higher in down-regulated genes relative to up-regulated genes (60% vs. 51%, respectively. P<0.05) (Figure 3D). The motifs enriched in down-regulated genes with NFIC peaks [-1000;TSS] include NFI, FOXM1, bHLH, RBPJ, and ARID5A; those enriched in up-regulated genes included NFI, TLX, HNF1, TCF3, and CTCF (Figure 3E and F, respectively). Gene set enrichment analysis showed that NFIC-bound down-regulated genes were associated with acinar differentiation, protein metabolism (e.g. oxidative phosphorylation, protein export, ribosome, seleno amino acid and purine metabolism, among others). Exemplary genes include *Amy2a5, Bhlha15, Cel, Cela1, Cela2b, Cpa1, Nr0b2, Pnliprp1, Rpl10a* and *Rpl23a* (Figure 3G). In contrast, NFIC-bound up-regulated genes were enriched in cell adhesion and inflammatory pathways (e.g. chemokine signaling, leukocyte transendothelial migration, ECM receptor interaction), MAPK signaling, and pathways in cancer including inflammatory genes (Figure 3H). Exemplary genes include *Cfi, Fos*, genes involved in ER stress and UPR *Dnajc5, Dnajc13, Hsp90aa1*, circadian clock regulator *Per2*, and *Pparg, Rara*, and *Rarg* (Figure 3H). These pathways have been shown to be critically relevant in pancreatic homeostasis and disease^23,27-32^. Representative examples of ChIP-Seq findings are displayed in Figure 3I.

### NFIC distinctly regulates the ribosomal program and aspects of the unfolded protein and ER stress responses

A striking finding from the GSEA analysis was the enrichment in gene sets related to protein synthesis (Supplementary Figure 6A). Multiple transcripts coding for ribosomal proteins were down-regulated in *Nfic*^-/-^ pancreata (Supplementary Figure 6B) and reactivity with an antibody detecting the 5.8S rRNA - a surrogate readout of ribosomes - was reduced in *Nfic*^-/-^ acinar cells (Figure 4A,B). In addition, there was a down-regulation of ER and Golgi complex components, including *Fkbp2, Dio* and *Pink1* that were bound by NFIC at their promoter region (Supplementary Figure 6C-E). The dysregulation of protein metabolism suggests a role of the mTOR pathway^33^. Western blotting analysis of *Nfic*^-/-^ pancreata showed reduced expression of phosphorylated ribosomal S6 Kinase 1 (P-RS6K1), its substrate S6 ribosomal protein (P-RS6), and phospho-elongation initiation factor 4e (P-EIF4E), together with a modest up-regulation of P-ERK (Figure 4C). IHC confirmed that these changes occur in acinar cells (Figure 4D-F). Interestingly, several ribosomal genes were found to be bound by NFIC in Chip-Seq experiments with a variety of cell types from the ENCODE project (Supplementary Figure 6F). These results indicate that NFIC loss impacts on ribosomal biogenesis and on the activity of the mTOR pathway in the adult mouse pancreas.

**Figure 6.**
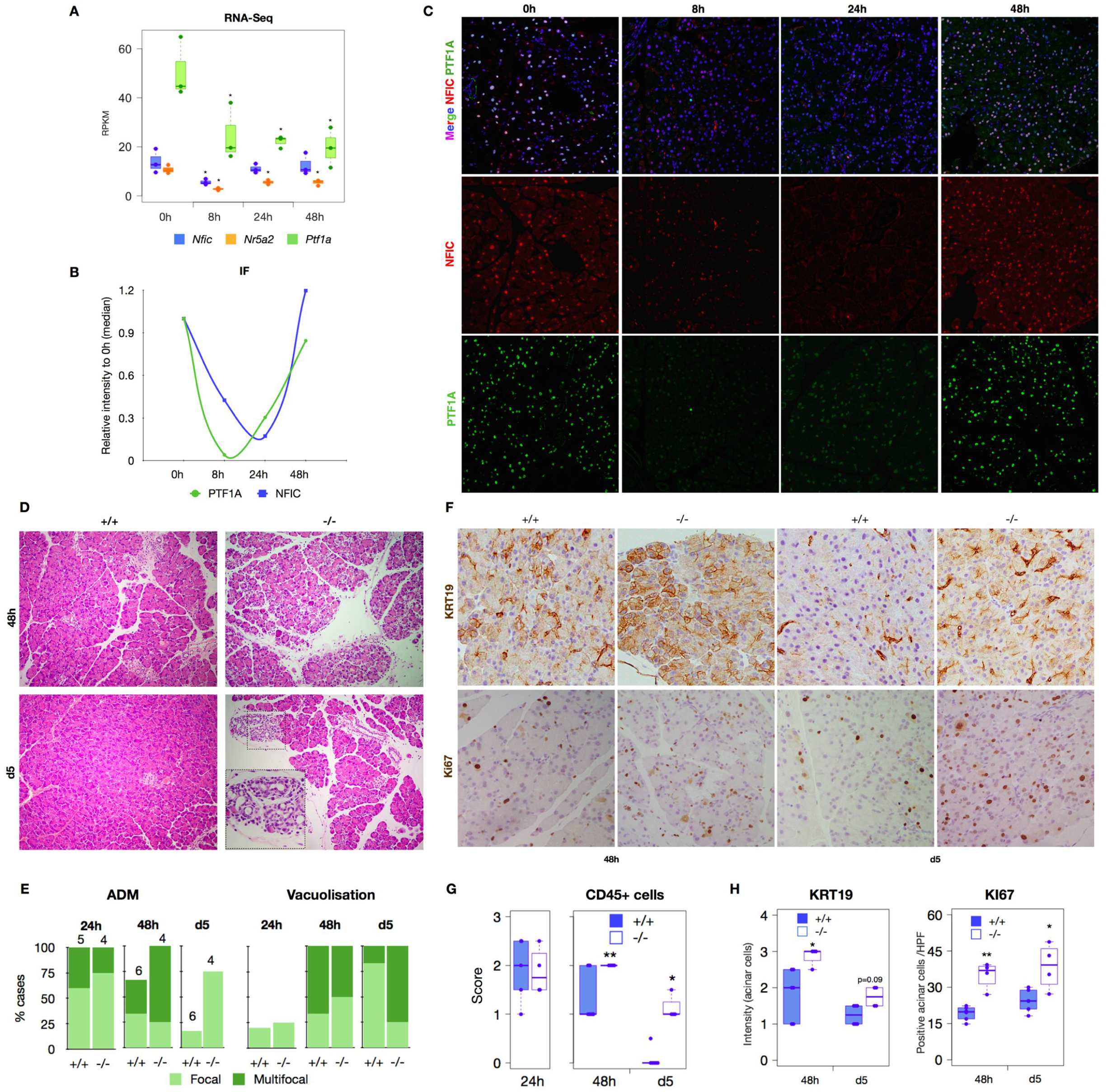
NFIC is dynamically regulated and required for a homeostatic response during caerulein pancreatitis. (A) RNA-Seq analysis of *Nfic, Nr5a2*, and *Ptf1a* expression in wild-type mice upon induction of a mild acute pancreatitis (n=3/group). Significance was calculated compared to expression at 0h. (B) IF analysis of NFIC and PTF1A upon pancreatitis induction showing NFIC down-regulation (n=4/group). (C) Quantification of PTF1A^+^ and NFIC^+^ cells in wild-type mice during pancreatitis. (D) Histological analysis of wild-type and *Nfic*^-/-^ pancreata 24h, 48h, and 5 days after the induction of pancreatitis showing increased damage in mutant mice. (E) Pancreatitis scoring indicates more severe damage in *Nfic*^-/-^ pancreata at 48h and day 5 (n>4/condition). (F-H) IHC reveals increased expression of KRT19, a higher number of KI67^+^ acinar cells, and increased infiltration by CD45^+^ cells in *Nfic*^-/-^ pancreata. Representative images (F). Quantification of CD45 (G), KRT19 and Ki67 expression (H). was subjectively scored (0-3) in *Nfic*-/-pancreata (n<u>=</u>4/group).

To determine whether similar findings apply to normal human pancreas, we used the GTEX dataset (n=171) and compared gene expression in samples with high vs. low *NFIC* levels (top vs. bottom 10 individuals): there was a −2.06-fold log2 difference in NFIC transcript levels in in *NFIC*^low^ vs. *NFIC*^high^ pancreata. Ninety-four percent of genes that were down-regulated in *Nfic*^-/-^ pancreata were also down-regulated in *NFIC*^low^ human pancreata (P=1.79 e^-47^) compared to 63% of a random gene list (Figure 4G). Among the common down-regulated genes are several involved in ribosomal function (Figure 4H), including *RPS5, RPS8, RPS11, RPS15, RPS21, RPS26*, and *RPS29* (Figure 4I). By contrast, only 27% of transcripts with up-regulated expression in *Nfic*^-/-^ pancreata were up-regulated in *NFIC*^low^ human samples, compared to 37% of a random list of genes (P=3.47e^-8^), including *RPS5, RPS8, RPS11, RPS15, RPS21, RPS26, RPS29* (Figure 4H,I). These data support a conservation of the function of NFIC in normal pancreas in mice and humans.

A large number of NFIC-bound down-regulated genes are involved in autophagy (e.g. *Ulk1, Prkaa2, Pik3c3, Gabarap, Gabarapl1, Map1lc3b, Sqstm1, Pink1, Dap*). Alterations in the unfolded protein response (UPR) and autophagy induce ER stress^34^. Accordingly, we found an up-regulation of transcripts of NFIC-bound genes coding for ER stress proteins (Figure 5A, Supplementary Figure 7A). Up-regulation of *Chop/Ddit3, Hspa5/Bip-1*, and spliced *Xbp1* (*sXbp1*) mRNAs was confirmed using RT-qPCR (Figure 5B). We observed a modest, significant, up-regulation of BIP-1 and CHOP in *Nfic*^-/-^ pancreata (Figure 5C,D); BIP-1 up-regulation was confirmed by IF (Supplementary Figure 7B,C). In addition, we found up-regulation of UPR genes in the pancreas of individuals with low *NFIC* including *HSPA90AA1, CALR3, HSPA6* and, to a lower extent, *CHOP, LDLR* and *TSEN15* (Supplementary Figure 7D) Moreover, NFIC binds to the proximal promoter or distal region of 28.31% (43/81) and 24.69% (20/81), respectively, of the genes associated to ER stress (Supplementary Figure 7E) (e.g. *Ddit4* and *Slc1a5*) (Supplementary Figure 7F). Interestingly, NFIC - but not NR5A2 - bound to the promoter of *Hspa5/Bip-1, Ddit3/Chop*, and *Hsp90aa1* (Figure 5E), highlighting that NFIC selectively regulates an aspect of the acinar secretory program related to the ER stress response.

**Figure 7.**
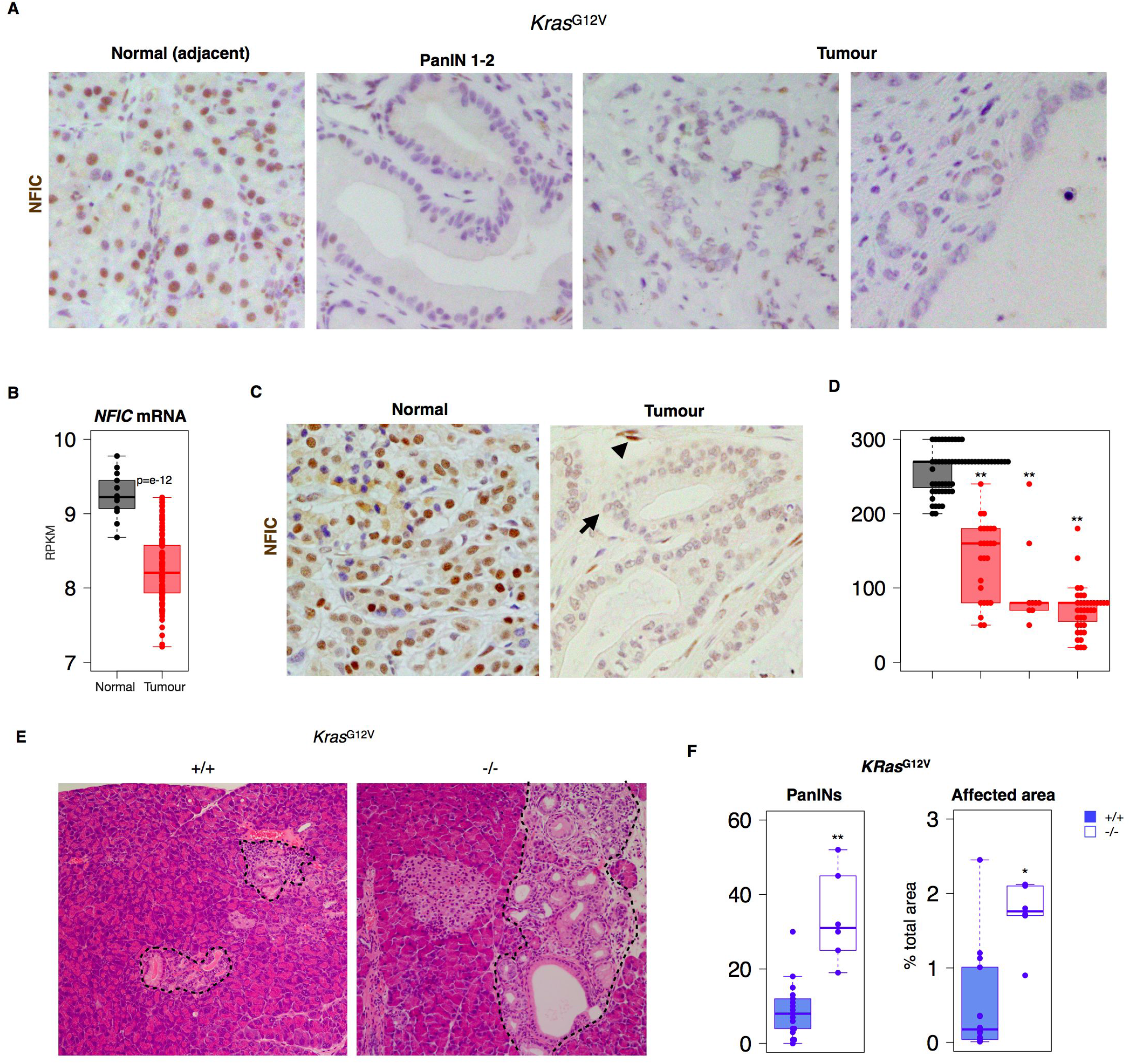
NFIC restrains formation of preneoplastic lesions in the pancreas. (A) IHC analysis of NFIC expression shows down-regulation in PanINs (right) and tumor cells compared to adjacent normal acinar cells (left). (B) *NFIC* mRNA analysis of tumor samples and normal adjacent tissue assessed by microarrays (Janky et al., 2016) showing reduced expression in tumor samples. (C,D) IHC analysis of NFIC expression in human PDAC specimens showing reduced expression in tumoral cells (arrow) compared to normal adjacent tissue or stromal cells (arrowheads). (E,F) Histological analysis of the pancreas of *KRas*^*G12V*^ or *KRas*^*G12V*^;*Nfic*^*-/-*^ 14-20 week old mice showing increased number of PanINs and area occupied by pre-neoplastic lesions (n>5/genotype).

To determine whether NFIC is involved in the ER stress response in the pancreas, 266-6 cells were treated with tunicamycin (TM), a protein N-glycosylation inhibitor. We also analyzed NR5A2 since it has been shown to participate in this process in the liver^35^. As expected, we observed a dose-dependent up-regulation of BIP-1 and a down-regulation of NFIC, NR5A2, and P-S6 at 24h. By 36h, NFIC levels remained low whereas NR5A2 expression had recovered (Supplementary Figure 7G). *Nfic* knock-down in 266-6 cells did not affect basal BIP-1 or CHOP expression but it sensitized cells to the effects of TM (Figure 5F,G). Accordingly, NFIC-overexpressing cells showed reduced expression of BIP-1 and CHOP upon treatment with TM (Figure 5H,I). In both cases, NR5A2 expression was unaffected (Figure 5F,H). Overall, these results indicate that NFIC regulates multiple aspects of protein synthesis biology and the ER stress response in pancreatic acinar cells.

### *Nfic* is required for recovery after induction of pancreatic damage

Acute pancreatitis is associated with a down-regulation of TFs involved in acinar differentiation and the up-regulation of ER stress and the UPR^36,37^. After induction of an acute caerulein pancreatitis (7 hourly doses) in wild type mice, *Nfic* mRNA levels decreased at early time points (8h) and were gradually restored upon recovery (Figure 6A). A similar expression pattern was observed at the protein level (Figure 6B,C). To investigate whether *Nfic* inactivation affects damage and/or regeneration, we induced a mild acute caerulein pancreatitis in control and knockout mice. At early time points (1-24h), wild-type and *Nfic*^-/-^ pancreata showed similar damage (Figure 6D). However, at 48h *Nfic*^-/-^ pancreata showed more prominent oedema, leukocyte infiltration, multifocal ADM, and acinar vacuolization. These differences persisted up to day 5 (Figure 6D,E and Supplementary Figure 7H). IHC confirmed an increased number of CD45^+^, KRT19^+^, and KI67^+^ acinar cells in *Nfic*^-/-^ pancreata (Figure 6F,G). These changes were accompanied by up-regulation of *Ddit3/Chop* and *Hsp17b11* mRNA in *Nfic*^-/-^ pancreata at 48h and day 5 (Figure 6H). These results indicate that NFIC is required for the recovery from pancreatic damage.

### NFIC suppresses PDAC initiation

Pancreatitis sensitizes the pancreas to the oncogenic effects of mutant *Kras*. We first analyzed NFIC expression in murine PanINs and PDAC from *Ptf1a-*Cre+/KI;*KRas*G12V+/KI (KC) using IHC and found that it is down-regulated in both preneoplastic and tumor cells (Figure 7A,B). Similar findings were made in samples from patients: we found significant down-regulation of *NFIC* mRNA in PDAC (n=118) when compared to normal tissue (n=13) (Figure 7B)^38^. Using IHC, we found that NFIC is consistently down-regulated in PanINs of low (n=56) and high (n=34) grade and in a subset of PDAC samples (n=43) (Figure 7C,D, right panel). Analysis of the PanCuRx microdissected PDAC dataset showed that NFIC mRNA expression was similar in classical- and basal-type tumors (not shown).

TFs involved in acinar differentiation have been shown to suppress tumor initiation in mice. Inactivation of *Nfic* in the context of the *Ptf1a-Cre+/KI;KRas*G12V+/KI alleles resulted in an increased number of PanINs (and of the relative area ocupied by them) in 18-24 week-old mice (Figure 7E,F). Altogether, these findings support a role of NFIC in the suppression of PDAC initiation.

## DISCUSSION

### NFIC is a novel regulator of the pancreatic acinar program

Acinar differentiation was long thought to be a “digital” process controlled by PTF1. Increasing evidence supports an “analog” differentiation model whereby additional TFs are required for “completion” of this process. Among them are NR5A2^6,7^, HNF1A^36^, GATA6^4^, MIST1^5,26^, and XBP1^39^. Here, we show that NFIC, a ubiquitous TF, is a novel acinar regulator present in a complex with NR5A2. While NFIC is not crucially required for pancreas organogenesis, in adult mice it regulates the ribosomal program and ER stress response and is dysregulated in pancreatitis and cancer.

We identified NFIC as an NR5A2 partner through co-binding in normal mouse pancreas but it remains to be determined whether both proteins interact directly. The conservation of spacing between NR5A2 and NFIC motifs among NR5A2 target genes supports transcriptional cooperation. However, comparison of NR5A2 ChIP-Seq data from embryonic and adult pancreas indicates that the role of NFIC is mainly in the latter, supporting its requirement for completion of acinar maturation and highlighting a functional role distinct from that of NR5A2. The transcriptional program driven by NFIC overlaps partially with that of the tissue-restricted PTF1A, NR5A2, and MIST1 factors, indicating that multiple TFs cooperate to activate acinar differentiation. However, inactivation of *Nfic* has milder effects than inactivation of *Ptf1a* or *Nr5a2*, possibly because the latter act at earlier stages of pancreatic development. NFI proteins were first proposed to be involved in the regulation of ubiquitous genes but they can also regulate tissue-specific genes^13^, including *CEL* in the mammary gland and *DSPP* in odontoblasts^14,19^. We show that NFIC also regulates the acinar program in the pancreas. The down-regulation of *CDH1* mRNA and protein observed in *Nfic*^-/-^ pancreata extends previous reports on *CDH1* regulation by NFIC in epithelial tissues^19,40^ and multiple aspects of cell adhesion were revealed by GSEA in the RNA-Seq analysis. *Nfic*^-/-^ pancreata also showed increased acinar proliferation and infiltrating leukocytes, associated with an up-regulation of inflammatory transcripts. This phenotype is similar to that of *Nr5a2*^+/-23^, *Hnf1a*^-/-36^ and not shown), *Gata6*^-/-4^, and *Ptf1a*^-/-25^ pancreata. The promoters of genes whose expression is up-regulated in *Nfic*^-/-^ pancreata were enriched in motifs for NF-kB, PPARy:RXRA, REL, and NFIC itself. These findings suggest that the activation of pro-inflammatory phenotypes in mice in which acinar cells fail to acquire normal maturation can result from both direct (NR5A2 and GATA6)^4,7,10,12^ and indirect (PTF1A and NFIC) mechanisms, in agreement with the ChIP-Seq data available.

### NFIC regulates expression of ribosomal genes and mitigates ER stress in the pancreas

A hallmark of acinar cells is their prominent capacity for protein synthesis, processing, and secretion^41^. This is achieved through acinar-specific transcriptional programs such as those driven by PTF1A^25^, MIST1^26^, and XBP1^39^ and - as shown here-NFIC. Accordingly, a coordinated down-regulation of gene sets related to the digestive process and to protein metabolism and oxidative phosphorylation occurs in adult *Nfic*^*-/-*^ pancreata. Similar changes are present in normal human pancreata displaying low *NFIC* expression, indicating the relevance to humans. The mTOR pathway is a central actor in protein biosynthesis and autophagy and, therefore, a candidate mediator of this phenotype (reviewed in [33]). We found reduced levels of P-RS6K1, its substrate P-RS6, and P-EIF4E, together with an up-regulation of P-ERK, in acinar cells of *Nfic*^*-/-*^ mice. However, a mechanistic link between NFIC and these signaling pathways is lacking.

The high level of basal protein synthesis in acinar cells underlies constitutive activation of the UPR to reduce ER stress^37,42^. We observed a down-regulation of UPR gene sets and an up-regulation of classical ER stress regulators in *Nfic*^*-/-*^ pancreata. The finding that NFIC - but not NR5A2 - binds the promoter of ER stress genes suggests a distinct role of the former in this process. This is supported by TM-mediated ER stress in 266-6 cells manipulated for NFIC gain-of-function and loss-of-function. We thus conclude that NFIC mitigates ER stress in acinar cells. Previous work has shown that NR5A2 is required for proper ER stress response in hepatocytes^35^. A re-analysis of published data shows that *Nfic* is down-regulated in *Nr5a2*^*-/-*^ hepatocytes in basal conditions (fold change of −0.54)^35^, suggesting that the deficient response to TM in *Nr5a2*^*-/-*^ hepatocytes might partially occur through *Nfic* down-regulation. Similarly, we found that NFIC is down-regulated in the pancreas upon TM-mediated ER stress (not shown), as has been reported in immortalized B cells^43^. These findings suggest that NFIC might be a broad regulator of ER stress response.

### NFIC is dynamically regulated during pancreatitis and cancer

A failure to achieve complete acinar maturation is associated with more severe damage and delayed recovery during caerulein-mediated pancreatitis, as shown upon inactivation of *Gata6, Mist1*, and *Ptf1a* in the pancreas^4,44^. In addition, disruption of the UPR and ER stress responses induces acinar damage and can lead to acute or chronic pancreatitis^45^. We found sustained up-regulation of *Ddit3/Chop* and *Hsp17b1* in *Nfic*^*-/-*^ pancreata upon induction of a mild pancreatitis indicating enhanced ER stress in the absence of *Nfic*.

Several groups have shown that mild defects in the regulation of pancreatic transcriptional programs can sensitize to pancreatitis and that TFs act as tumor suppressors^10,11,12^. The role of NFIC in acinar cell differentiation and mitigation of ER stress suggested a contribution during tumorigenesis. NFIC has been proposed to act as a tumor suppressor in breast cancer, as it activates *TP53*, represses *CCND1* and *FOXF1*, and is down-regulated by *c*-*MYC* and *Ha*-*RAS* oncogenes^15-17,46,47^. In breast cancer, NFIC is down-regulated and high expression is associated with better prognosis^16^. Deregulation of other NFI family members has been reported in several tumor types: NFIB is overexpressed in metastatic neuroendocrine lung tumors and it drives metastatic progression of small cell lung cancers by increasing chromatin accessibility^48,49^. The lack of *NFIC* mutations and/ or genomic alterations in human PDAC (https://cancergenome.nih.gov/newsevents/newsannouncements/pancreatic_2017) suggests that other mechanisms may contribute to tumor development/progression.

We now show that NFIC is a novel, ubiquitous, TF with tissue-specific functions in the pancreas that cooperates with NR5A2 to binds target genes and controls their expression *in vitro* and *in vivo*. Unlike other pancreatic TFs previously described, the role of NFIC is restricted to the adult pancreas and distinctly affects RNA and protein metabolism and the UPR-mediated ER stress. Mutations leading to protein misfolding, the UPR, and activation of ER stress cause chronic pancreatitis and can contribute to the risk of PDAC^50^, further supporting the role of NFIC in pancreatic homeostasis and disease.

## MATERIAL AND METHODS

### Mice and experimental manipulations

The following mouse strains were used: *Nfic*^-/-18^, *Ptf1a*^+/Cre^ knock-in^51^, and *KRas*^G12V^ conditional knock-in ^8^. All crosses were maintained in a predominant C57BL/6 background. Experiments were performed using 8-12 week-old mice of both sexes, except for glucose tolerance tests where only males were used. Littermate mice were used as controls. All animal procedures were approved by local and regional ethics committees (Institutional Animal Care and Use Committee and Ethics Committee for Research and Animal Welfare, Instituto de Salud Carlos III) and performed according to the European Union guidelines.

A mild acute pancreatitis was induced by 7 hourly injections of the cholecystokinin analogue caerulein (Bachem) at 50 μg/kg. In brief, animals were weighed before the procedure and caerulein was administered intraperitoneally. Mice were killed by cervical dislocation after 1 & 4h after the last caerulein injection or 24, 48h and 5 days after the first caerulein injection. For the glucose tolerance test, male mice were fasted for 16 h and basal glycaemia was measured in tail blood. Mice received a glucose solution (2g/kg) administered intraperitoneally and glycaemia was measured 15, 30, 60, and 120 min later using an automated glucose monitor (Accu-ChekⓅAviva). Fasting glucose was considered as baseline (0h). The number of mice used in each experiment is shown in the legend of each figure. For most experiments, ≥5 mice per group were used. No specific randomization method was used.

### Acinar cell isolation

Acinar cells were isolated by collagenase P (1mg/pancreas) digestion and maintained at 37 °C for 24 h in RPMI containing L-glutamine, 1mM of pyruvate (Sigma-Aldrich), soybean trypsin inhibitor (STI) (Gibco, 17075-029) (0.1mg/ml) and 10% foetal bovine serum. (ref 23)

### Histology, immunofluorescence (IF) and immunohistochemical (IHC) analyses

Pancreata were immediately placed in buffered formalin or 4% paraformaldehyde. Histological processing was performed using standard procedures. To score damage in acute pancreatitis experiments, inflammation-related histological parameters (oedema, inflammatory cell infiltration, vacuolization, and acino-ductal metaplasia [ADM]) were scored blindly by IC and FXR according to the grade of severity (0-3).

IF and IHC analyses were performed using 3 μm sections of formalin-fixed paraffin-embedded tissues, unless otherwise indicated. After deparaffinization and rehydration, antigen retrieval was performed by boiling in citrate buffer pH 6 for 10 min. For IF, sections were incubated for 45 min at room temperature with 3% BSA, 0.1% Triton X-100-PBS and then with the primary antibody overnight at 4 °C. For double or triple IF, the corresponding antibodies were added simultaneously and incubated overnight at 4 °C. Sections were then washed with 0.1% Triton–PBS, incubated with the appropriate fluorochrome-conjugated secondary antibody, and nuclei were counter-stained with DAPI. After washing with PBS, sections were mounted with Prolong Gold Antifade Reagent (Life Technology).

For IHC analyses, after antigen retrieval, endogenous peroxidase was inactivated with 3% H_2_O_2_ in methanol for 30 min at room temperature. Sections were incubated with 2% BSA-PBS for 1h at room temperature, and then with the primary antibody overnight at 4 °C. After washing, the Envision secondary reagent (DAKO) was added for 40 min at room temperature and sections were washed x3 with PBS. 3,30-Diaminobenzidine tetrahydrochloride (DAB) was used as a chromogen. Sections were lightly counterstained with haematoxylin, dehydrated, and mounted. For some antibodies, an automated immunostaining platform was used (Ventana Discovery XT, Roche). A non-related IgG was used as a negative control. To validate the specificity of anti-NFIC antibodies, *Nfic*^-/-^ pancreata were used as controls.

For CD45 quantification, whole digital slide images were acquired with an Axio Scan Z1, Zeiss scanner and then captured with the Zen Software (Zeiss). Image analysis and quantification were performed with the AxioVision software package (Zeiss). Briefly, areas of interest (AOI) were selected for quantification and then exported as individual TIFF images. CD45 staining were quantified using AxioVision 4.6 (Zeiss). Data obtained were then compiled and appropriately assessed. Images containing lymph nodes, and with artifactual staining or suboptimal cutting were eliminated from the analysis.

For quantification of KI67^+^ positive cells, at least 10 random images from each pancreas were selected and only positive acinar cells were quantified. For BIP-1 quantification, at least 10 random images from each pancreas were taken and fluorescence intensity was calculated using FIJI software (https://fiji.sc/). For semi-quantitative analysis of KRT19 staining, intensity was scored from 0-3 by IC.

A list of antibodies used for IHC and IF is provided in Supplementary Table 3.

### Quantitative RT-PCR (RT-qPCR)

For RNA isolation, pancreata were homogenized in denaturing buffer (4 M guanidine thiocyanate, 0.1 M Trizma HCl pH 7.5, 1% 2-mercaptoethanol) and processed as described earlier^23^. Total RNA was treated with DNase I (Ambion) for 30 min at 37 °C and cDNAs were prepared according to the manufacturer’s specifications, using the TaqMan reverse transcription reagents (Applied Biosystems, Roche). qRT-PCR analysis was performed using the SYBR Green PCR master mix and an ABIPRISM 7900HT instrument (Applied Biosystems). Expression levels were normalized to endogenous *Hprt* mRNA levels using the ΔΔ*C*_*t*_ method. The results shown are representative of at least four biological replicates. The sequence of the primers used is provided in Supplementary Table 4.

### Immunoprecipitation and western blotting

Pancreata were snap-frozen for protein isolation. For immunoprecipitation of proteins from fresh total pancreas lysates, a piece of mouse pancreas was isolated and minced in 50 mM Tris-HCl pH 8, 150 mM NaCl, 5 mM EDTA, 0.5% NP-40 containing 3× phosphatase inhibitor cocktail (Sigma-Aldrich) and 3× EDTA-free complete protease inhibitor cocktail (Roche). Lysates were briefly sonicated until the protein solution was clear, cleared for 10 min at 11,000 rpm at 4 °C and the supernatant was recovered. Antibody-coated protein A or protein G dynabeads (Life Technology) were used for immunoprecipitation. In brief, beads were washed three times with PBS and incubated with anti-NR5A2 or normal goat IgG (Millipore) overnight at 4 °C. After washing three times with PBS and twice with coupling buffer (27.3 mM sodium tetraborate, 72.7 mM boric acid), the dry beads were incubated overnight at 4 °C in freshly prepared 38 mM dimethyl pimelimidate dihydrochloride in 0.1 M sodium tetraborate. Afterwards, beads were washed three times with coupling buffer and once with 1 M Tris pH 9. Then, 1 ml of the Tris solution was added to the beads and incubated for 10 min at room temperature with rotation to block amino groups and stop crosslinking. Finally, beads were washed three times with storage buffer (6.5 mM sodium tetraborate/boric acid) and stored at 4 °C until used. Protein lysates (10-15 mg, tissues) were then incubated overnight at 4 °C with antibody-coated dynabeads (Thermo Fisher Scientific). Bound immune complexes were washed twice with lysis buffer containing NP-40, and then eluted by boiling in 2× Laemmli buffer (10% glycerol, 2% sodium dodecyl sulphate and 0.125 M Tris-HCl pH 6.8) for 5 min.

For western blotting, proteins were extracted from pancreatic tissue, isolated acinar cells or cultured cells using either Laemmli buffer, lysis buffer (50 mM Tris-HCl pH 8, 150 mM NaCl, 5 mM EDTA and 0.5% NP-40) or 5M urea, supplemented with protease inhibitor and phosphatase inhibitor cocktails. Protein concentration was measured using the BCA reagent (Biorad), Nanodrop or extrapolated when using Laemmli lysis buffer. Proteins were resolved either by standard SDS-PAGE or 4-20% TGX pre-cast gels (Biorad) and transferred onto nitrocellulose membranes. A list of antibodies used for WB, ChIP and IP is provided in Supplementary Table 3. Densitometry analysis of digitalised western blotting images was performed using Fiji software (https://fiji.sc/).

### Chromatin immunoprecipitation (ChIP)

Pancreas tissue was minced, washed with cold PBS supplemented with 3× protease and phosphatase cocktail inhibitors, and then fixed with 1% formaldehyde for 20 min at room temperature. Glycine was added to a final concentration of 0.125 M for 5 min at room temperature. The fixed tissue was soaked in SDS buffer (50 mM Tris pH 8.1, 100 mM NaCl, 5 mM EDTA and 0.5% SDS) and homogenized using a douncer. The supernatant was collected after centrifugation and chromatin was sonicated with a Covaris instrument for 40 min (20% duty cycle; 10% intensity; 200 cycle), yielding DNA fragments with a bulk size of 300-500 bp. Samples were centrifuged to pellet cell debris. The amount of chromatin isolated was quantified using Nanodrop; an aliquot of this material was used as input for final quantification. Samples (0.5-1 mg of chromatin) were diluted with Triton buffer (100 mM Tris pH 8.6, 0.3% SDS, 1.7% Triton X-100 and 5 mM EDTA) to 1ml and pre-cleared for 2 h with a mix of protein A and G (previously blocked with 5% BSA) at 4 °C. Antibody-coated beads were added: anti-NR5A2 (2 μg), anti-NFIC (1 μg), and rabbit anti-PTF1A serum (1/500). Non-related IgG was used as a control. After incubating for 3 h at 4 °C in a rotating platform, beads were successively washed with 1 ml of mixed micelle buffer (20 mM Tris pH 8.1, 150 mM NaCl, 5 mM EDTA, 5% w/v sucrose, 1% Triton X-100 and 0.2% SDS), buffer 500 (50 mM HEPES at pH 7.5, 0.1% w/v deoxycholic acid, 1% Triton X-100, 500 mM NaCl and 1 mM EDTA), LiCl detergent wash buffer (10 mM Tris at pH 8.0, 0.5% deoxycholic acid, 0.5% NP-40, 250 mM LiCl and 1 mM EDTA) and TE (pH 7.5), and then bound molecules were eluted by incubating overnight in elution buffer (containing 1% SDS and 100 mM NaHCO_3_) at 65 °C, and treated with proteinase K solution (10 M EDTA, 40 mM Tris-HCl pH 6.5, 40 μg/ml proteinase K). The eluted DNA was purified by phenol– chloroform extraction. After isolation, pelleted DNA was resuspended in nuclease-free water (150 μl). Gene occupancy was then analysed by real-time PCR using 1 μl of the eluted DNA diluted in a final volume of 10 μl. The sequence of the primers used for ChIP-qPCR is provided in Supplementary Table 4.

### ChIP-Seq

ChIP sequencing libraries were prepared from purified DNA using “NEBNext Ultra II DNA Library Prep Kit for Illumina” from New England BioLabs (E7645), as per the manufacturers instructions. The resulting libraries were sequenced on Illumina HiSeq 2500, v4 Chemistry.

### NFIC knockdown

NFIC expression was interfered in 266-6 cells using Mission shRNA lentiviral constructs purchased from Sigma-Aldrich. *Nfic* sh1 [TRCN0000374154 targeting ACAGACAGCCTCCACCTACTT), *Nfic* sh2 (TRCN0000310992 targeting TGTGTGCAGCCGCACCATATT), and *Nfic* sh3 (TRCN0000301779, targeting GATGGACAAATCTCCATTCAA)]. Control cells were transformed using lentiviral particles transducing the scrambled vector CCGGCAACAAGATGAAGAGCACCAACTCGAGTTGGTGCTCTTCATCTTGTTGTTTTT (shNT).

To produce lentiviral particles, HEK293-FT cells (ATCC) were allowed to reach 50% of confluence and transfected with 15 μg of shNT, *Nfic* sh1, *Nfic* sh2 or *Nfic sh3* plasmids together with 8 μg of psPAX and 2 μg of pCMV-VSVG helper plasmids using CaCl_2_ 2M HBSS. After 12 h, the supernatant was collected and replaced with 5 ml of fresh medium. The supernatant was collected 24h, 48 h and 72 h after transfection. The medium was filtered (0.45 μm pore) and added to 266-6 cells (at 50–60% of confluence); 1 µg/mL of Polybrene (hexadimethrine bromide, Sigma-Aldrich 107689) was added to increase infection efficiency. After 2-3 rounds of infection, the supernatant was removed and replaced with fresh medium. One day later, puromycin (1-2 μg/ml) (Sigma-Aldrich) was added and two days later, the medium was replaced.

### NFIC lentiviral overexpression

*Nfic-*HA tagged cDNA was purchased from Addgene (https://www.addgene.org/31403/) and subcloned into the lentiviral vector pLVX-puro using *XhoI* and *XbaI*. Insert sequence was checked using enzymatic digestion and Sanger sequencing. The production of lentiviral particles and cellular infection were performed as described earlier. The medium from the transfectants was collected 24h, 48h and 72h after transfection. Subsequently, 266-6 cells were infected using Polybrene as described earlier. After selection with puromycin for 24-48h, resistant 266-6 cells were collected for RNA and protein analysis.

### Tunicamycin (TM) treatment

266-6 cells were seeded until they reached 70% confluence. After pilot dose-response experiments, a concentration of 10nM was chosen; cells treated with TM or vehicle were collected at various time-points for RNA and protein analysis.

### RNA-Seq libraries preparation and analysis

RNA from wild type and Nfic-/-pancreata was isolated as described above and sequenced on Illumina platform. RNA-seq data for Nr5a2 (GSE34030), Mist1 (GSE86288) were downloaded from SRA. Data were analysed using the nextpresso pipeline http://bioinfo.cnio.es/nextpresso/).

Comparison of gene expression in normal human pancreata according to NFIC transcript levels was performed using RNA-Seq data from GTEX website (https://gtexportal.org/home/datasets, version 6) (n=171), as described21. The expression data matrix was sorted by NFIC expression levels taking the 10 individuals scoring highest and lowest NFIC expression levels (NFIChigh, NFIClow). Differential expression analysis using the DEGseq package of R (https://bioconductor.org/packages/release/bioc/html/DEGseq.html). MA-plot-based method with Random Sampling model -MARS-(Wang et al. 2009) was applied and only genes with significance P<0.001 were used in the analysis. All data have been deposited in GEO with accession number GSE126907.

### RNA-seq and data processing

RNA-Seq of pancreata from wild type mice during pancreatitis was analysed as previously described in [23] and is available under GSE84659 (https://www.ncbi.nlm.nih.gov/geo/query/acc.cgi?acc=GSE84659).

Briefly, RNA from wild type and Nfic-/-pancreata was isolated as described above. Data were analysed using the nextpresso pipeline http://bioinfo.cnio.es/nextpresso/). Tophat was used for alignment (tophat-2.0.10.Linux_x86_64) using the following parameters: useGTF=“true” nTophatThreads=“1” maxMultihits=“5” readMismatches=“1”, segmentLength=“19”, segmentMismatches=“1”, spliceMismatches=“0”, reportSecondaryAlignments=“false” bowtie=“1”, readEditDist=“2” readGapLength=“2” referenceIndexing=“false”, --no-coverage-search. Gene expression was quantified using cufflinks (version 2.2.1) using the following parameters: useGTF=“true” nThreads=“1” fragBiasCorrect= “true”, multiReadCorrect=“false” library Normalization Method= “classic-fpkm” max Bundle Frags=“5000000000”, normalization= “compatibleHits”, no Effective Length Correction=“true” no Length Correction=“false”. Differential expression analysis was done using cuffdiff (version 2.2.1) using the following parameters: useCuffmergeAssembly=“false”, nThreads=“1”, fragBiasCorrect=“true”, multiReadCorrect=“false”, libraryNormalization Method=“geometric” FDR=“0.05” minAlignmentCount=“10”, seed=“123L” FPKMthreshold=“2”, maxBundleFrags= “5000000000”, noEffectiveLengthCorrection=“true”, noLengthCorrection=“false” dispersion Method= “pooled”. Normalised expression across all samples was calculated using cuffnorm (vs 2.2.1) using the following parameters: useCuffmergeAssembly=“false”, nThreads=“1”, output Format=“simple-table” libraryNormalizationMethod=“geometric”, seed=“123L” normalization= “compatibleHits”. BEDTools-Version-2.16.2 and samtools-0.1.19; bowtie-1.0.0 were also used to execute the software shown above.

To analyze gene expression in normal human pancreas, RNA-Seq data was downloaded from GTEX website (https://gtexportal.org/home/datasets, version 6); 171 samples were used. The expression data matrix was sorted by NFIC expression levels to then take the 10 top and bottom individuals of expression (NFIChigh, NFIClow) top vs. bottom 10 individuals. Differential expression analysis using the DEGseq package of R (https://bioconductor.org/packages/release/bioc/html/DEGseq.html). MA-plot-based method with Random Sampling model -MARS-(Wang et al. 2009) was applied and only genes with significance P<0.001 were used in the analysis.

### Principal component analysis (PCA)

The Pearson correlation was calculated from the expression value (expressed as fragments per kilobase of transcript per million mapped reads) of each gene for each sample by using the ‘cor’ command in R (https://www.r-project.org/). Principal component analysis was performed using the ‘prcomp’ command in R, from the correlation value of each sample.

### Gene Set Enrichment Analysis (GSEA)

A ranking metric [-log10(p value)/sign(log2FoldChange)] was used to generate a ranked gene list from the DEseq output. The list of pre-ranked genes was then analysed with using the molecular signature dataset of GSEA for Gene Ontology (GO), KEGG, REACTOME, HALLMARKS or CANONICAL PATHWAYS databases as described in the Figure legends and the text. Significantly enriched terms were identified using a false discovery rate (FDR) *q* value of <0.25.

### NFIC ChIP-Seq

Chromatin from mouse pancreas tissue was extracted and processed as described above. For ChIP sequencing, libraries were prepared from purified DNA using “NEBNext Ultra II DNA Library Prep Kit for Illumina” from New England BioLabs (NEB, #E7645), as per the manufacturers’ instructions. The resulting libraries were sequenced on Illumina HiSeq 2500, v4 Chemistry.

### ChIP-seq data processing

Data from NR5A2 ChIP-Seq in adult pancreata (SRR389293, SRR389294), NR5A2 ChIP-Seq in ES cells (GSM470523, GSM470524), PTF1A ChIP-Seq in adult pancreata (GSM2051452, GSM2051453), and MIST1 ChIP-Seq in adult pancreata (GSM2299654,GSM2299654, GSM2299655) were downloaded from the Gene Expression Omnibus website (https://www.ncbi.nlm.nih.gov/geo/) and analysis was performed as described^23^. Briefly, after the quality check by fastqc (v.0.9.4, Babraham Bioinformatics), the alignment and peak calling for the ChIP-seq data was performed using RUbioSeq+ pipeline^52^. Merging of replicate peaks and peak annotation was done using HOMER. Peak calling, annotation and motif enrichment was identified using HOMER (Heinz et al., 2010; http://homer.ucsd.edu/homer/). Reads were directionally extended to 300 bp and, for each base pair in the genome, the number of overlapping sequence reads was determined and averaged over a 10-bp window to create a wig file to visualize the data in the University of California Santa Cruz (UCSC) genome browser.

NFIC ChIP-Seq data using GM12878, ECC1, HepG2, SK-N-SH and K562 cells were downloaded from (https://www.encodeproject.org/targets/NFIC-human/). ChIP-Seq peaks were analysed using Peak Analyser_1.4, using and Nearest Transcription Start Site parameter was used to annotate the genomic location of peaks. More than 95% of the target genes identified in the replicate with lowest number of target genes were included in the replicate with highest number. Among the two replicates, the one with highest number of identified target genes was taken: replicate 1 of NFIC ChIP-Seq in GM12878, NFIC ChIP-Seq in HepG2 and NFIC ChIP-Seq in SK-N-SH and replicate 2 of NFIC ChIP-Seq in ECC1 cell line.

### Other statistical analyses

Comparisons of quantitative data between groups were was performed using one-sided Mann-Whitney U test in all cases for which there was a prior hypothesis, except for the data shown in Figure 6 D, E where a prior existed. Box plots represent the median and second and third quartiles (interquartile range, IQR) of the data. Error bars are generated by R software and represent the highest and lowest data within 1.5× IQR range. All statistical analyses were performed with Excel, R software, https://ccb-compute2.cs.uni-saarland.de/wtest/ or https://www.medcalc.org/calc/comparison_of_proportions.php. The random list of genes were generated using https://www.dcode.fr/random-selection and http://www.molbiotools.com/randomgenesetgenerator.html websites. Duplicated transcripts in RNA-Seq data were deleted for analysis. Dotted line refer to threshold for statistical significance (-log10[0.25]=0.60) or (-log10[0.05]=1.30. Two-sided Mann-Whitney U test was used unless otherwise indicated; P<0.05 (*); P<0.01 (**)

## Abbreviations

ChIP: chromatin immunoprecipitation
DEG: differentially expressed genes
EMT: epithelial-mesenchymal transition
ER: endoplasmic reticulum
GSEA: Gene set enrichment analysis
IF: immunofluorescence
IHC: immunohistochemistry
PDAC: pancreatic ductal adenocarcinoma
TF: transcription factor
TM: tunicamycin
UPR: unfolded protein response

## Acknowledgements

We thank A. Efeyan, N. Djouder, and P. Martinelli for valuable discussions; M. Soengas and B. Bréant for providing antibodies; J. Perales and F. Al-Shahrour for help handling the GTEX dataset; R.M. Gronostajski, M. Barbacid, and C.W. Wright for providing mice; M. Barba, C. Yolanda and T. Lobato for technical assistance; and the Core Facilities and Bioinformatics Unit of Spanish National Cancer Research Center (CNIO) for support.

## Notes

<u>Conflicts of interest</u>: none to declare

<u>Funding</u>: This work was supported, in part, by grants SAF2011-29530, SAF2015-70553-R, and RTI2018-101071-B-I00 from Ministerio de Ciencia, Innovación y Universidades (Madrid, Spain) (co-funded by the ERDF-EU) and RTICC from Instituto de Salud Carlos III (RD12/0036/0034) to FXR. IC was recipient of a Beca de Formación del Personal Investigador from Ministerio de Economía y Competitividad (Madrid, Spain). The research leading to these results has received funding from People Programme (Marie Curie Actions) of the European Union’s Seventh Framework Programme (FP7/2007- 2013) (REA grant agreement n° 608765”). SP was supported by a Juan de la Cierva Programme fellowship from Ministerio de Ciencia, Innovación y Universidades. IM was supported by a Fellowship from Fundació Bancaria La Caixa (ID 100010434) (grant number LCF/BQ/ES18/11670009). CNIO is supported by Ministerio de Ciencia, Innovación y Universidades as a Centro de Excelencia Severo Ochoa SEV-2015-0510.

### Competing Interest Statement

The authors have declared no competing interest.

## REFERENCES

1. Pandol SJ. The exocrine pancreas. San Rafael (CA), Morgan & Claypool Life Sciences 2010.

2. Rose SD, Swift GH, et al. The role of PTF1-P48 in pancreatic acinar gene expression. J Biol Chem 2001; 276:44018–44026.

3. Beres TM, Masui T, et al. PTF1 is an organ-specific and Notch-independent basic helix–loop-helix complex containing the mammalian Suppressor of Hairless (RBP-J) or its paralogue, RBP-L. Mol Cell Biol 2006; 26:117–130.

4. Martinelli P, Cañamero M, et al. Gata6 is required for complete acinar differentiation and maintenance of the exocrine pancreas in adult mice. Gut 2013; 62:1481–1488.

5. Pin CL, Rukstalis JM, et al. The bHLH transcription factor Mist1 is required to maintain exocrine pancreas cell organization and acinar cell identity. J Cell Biol 2001; 155:519–530.

6. Holmstron SR, Deering T, et al. LRH-1 and PTF1-L coregulate an exocrine pancreas-specific transcriptional network for digestive function. Genes & Dev 2011; 25:1674–1679.

7. Hale MA, Swift GH, et al. Development. The nuclear hormone receptor family member NR5A2 controls aspects of multipotent progenitor cell formation and acinar differentiation during pancreatic organogenesis. Development 2014; 141:3123–3133.

8. Guerra C, Schuhmacher AJ, et al. Chronic pancreatitis is essential for induction of pancreatic ductal adenocarcinoma by K-Ras oncogenes in adult mice. Cancer Cell 2007; 11:291–302.

9. Kopp JL, von Figura G, et al. Identification of SOX9-dependent acinar-to-ductal reprogramming as the principal mechanism for initiation of pancreatic ductal adenocarcinoma. Cancer Cell 2012; 22:737–750.

10. Flandez M, Cendrowski J, et al. Nr5a2 heterozygosity sensitises to, and cooperates with, inflammation in KRas(G12V)-driven pancreatic tumourigenesis. Gut 2014; 63:647–655.

11. Martinelli P, Madriles F, et al. The acinar regulator Gata6 suppresses KrasG12V-driven pancreatic tumorigenesis in mice. Gut 2016; 65:476–486.

12. Von Figura G, Morris JP, et al. Nr5a2 maintains acinar cell differentiation and constrains oncogenic Kras-mediated pancreatic neoplastic initiation. Gut 2014; 63:656–664.

13. Gronostajski RM. Roles of the NFI/CTF gene family in transcription and development. Gene 2000; 249:31–45.

14. Kannius-Janson M, Johansson EM, et al. Nuclear factor 1-C2 contributes to the tissue-specific activation of a milk protein gene in the differentiating mammary gland. J Biol Chem 2002; 277: 17589–17596.

15. Eeckhoute J, Carroll JS, et al. Cell-type-specific transcriptional network required for estrogen regulation of cyclin D1 and cell cycle progression in breast cancer. Genes Dev 2006; 20:2513–2526.

16. Nilsson J, Helou K, et al. Nuclear Janus-activated kinase 2/nuclear factor 1-C2 suppresses tumorigenesis and epithelial-to-mesenchimal transition by repressing Forkhead box F1. Cancer Res 2010; 70:2020–2029.

17. Johansson EM, Kannius-Janson M, et al. The p53 tumor suppressor gene is regulated in vivo by nuclear factor 1-C2 in the mouse mammary gland during pregnancy. Oncogene 2003; 22:6061–6070.

18. Steele-Perkins G, Butz KG, et al. Essential role for NFI-C/CTF transcription-replication factor in tooth root development. Mol Cell Biol 2003; 23:1075–84.

19. Lee HK, Lee DS. Nuclear Factor I-C (NFIC) regulates Dentin Sialophosphoprotein (DSPP) and E-cadherin via control of Krüppel-like Factor 4 (KLF4) during dentinogenesis. J Biol Chem 2014; 289:28225–28236.

20. Zhou J, Wang S, et al. Nuclear factor I-C reciprocally regulates adipocyte and osteoblast differentiation via control of canonical Wnt signaling. FASEB J 2017; 31:1939–1952.

21. Liu Y, Feng J, et al. An Nfic-hedgehog signaling cascade regulates tooth root development. Development 2015; 142:3374–3382.

22. Heng JC, Feng B, et al. The nuclear receptor Nr5a2 can replace Oct4 in the reprogramming of murine somatic cells to pluripotent cells. Cell Stem Cell 2010; 6:167–174.

23. Cobo I, Martinelli P, et al. Transcriptional regulation by NR5A2 links differentiation and inflammation in the pancreas. Nature 2018; 554:533–537.

24. Cendrowski J, Lobo VJ, et al. Mnk1 is a novel acinar cell-specific kinase required for exocrine pancreatic secretion and response to pancreatitis in mice. Gut 2015: 64;937–947.

25. Hoang CQ, Hale MA, et al. Transcriptional Maintenance of Pancreatic Acinar Identity, Differentiation, and Homeostasis by PTF1A. Mol Cell Biol 2016; 36:3033–3047.

26. Jiang M, Azevedo-Pouly AC, et al. MIST1 and PTF1 Collaborate in Feed-Forward Regulatory Loops That Maintain the Pancreatic Acinar Phenotype in Adult Mice. Mol Cell Biol 2016. 36:2945–2955.

27. Biasci D, Smoragiewicz M, et al. CXCR4 inhibition in human pancreatic and colorectal cancers induces an integrated immune response. Proc Natl Acad Sci U S A. 2020.17;117:28960–28970.

28. Heinrich EL, Lee W, et al. Chemokine CXCL12 activates dual CXCR4 and CXCR7-mediated signaling pathways in pancreatic cancer cells. J Transl Med 2012;10: 68.

29. Perera RM, Bardeesy N. Pancreatic Cancer Metabolism: Breaking It Down to Build It Back Up. Cancer Discov. 2015;5:1247–1261.

30. Qin C, Yang G, et al. Metabolism of pancreatic cancer: paving the way to better anticancer strategies. Mol Cancer. 2020;19:50.

31. García-Costela M, Escudero-Feliú J, et al. Circadian Genes as Therapeutic Targets in Pancreatic Cancer. Front Endocrinol (Lausanne). 2020;11:638.

32. Kobayashi H, Spilde TL, et al. Retinoid signaling controls mouse pancreatic exocrine lineage selection through epithelial-mesenchymal interactions. Gastroenterology. 2002;123:1331–1340.

33. Laplante M and Sabatini DM. mTOR signaling in growth control and disease. Cell 2012; 149:274–293.

34. Senft D and Ronai ZA. UPR, autophagy, and mitochondria crosstalk underlies the ER stress response. Trends Biochem Sci 2015; 40:141–148.

35. Mamrosh JL, Lee JM, et al. Nuclear receptor LRH-1/NR5A2 is required and targetable for liver endoplasmic reticulum stress resolution. Elife 2014; 3:e01694.

36. Molero X, Vaquero EC, et al. Gene expression dynamics after murine pancreatitis unveils novel roles for Hnf1alpha in acinar cell homeostasis. Gut 2012; 61:1187–1196.

37. Kubisch CH, Logsdon CD. Secretagogues differentially activate endoplasmic reticulum stress responses in pancreatic acinar cells. Am J Physiol Gastrointest Liver Physiol 2007; 292:G1804–G1812.

38. Janky R, Binda MM, et al. Prognostic relevance of molecular subtypes and master regulators in pancreatic ductal adenocarcinoma. BMC Cancer 2016; 16:632.

39. Lee AH, Chu GC, et al. XBP-1 is required for biogenesis of cellular secretory machinery of exocrine glands. EMBO J 2005; 24:4368–4380.

40. Lee HK, Lee DS, et al. Nuclear factor I-C regulates E-cadherin via control of Klf4 in breast cancer. BMC Cancer 2015; 15:113.

41. Case RM. Synthesis, intracellular transport and discharge of exportable proteins in the pancreatic acinar cell and other cells. Biol Rev Camb Philos Soc 1978; 53:211–354.

42. Drummond A and Wilke CO. The evolutionary consequences of erroneous protein synthesis. Nat Rev Genet 2009; 10:715–724.

43. Dombroski BA, Nayak RR, et al. Gene expression and genetic variation in response to endoplasmic reticulum stress in human cells. Am J Hum Genet 2010; 86:719–729.

44. Kowalik AS, Johnson CL, et al. Mice lacking the transcription factor Mist1 exhibit an altered stress response and increased sensitivity to caerulein-induced pancreatitis. Am J Physiol Gastrointest Liver Physiol. 2007;292:G1123–1132.

45. Sah RP, Garg SK, et al. Endoplasmic reticulum stress is chronically activated in chronic pancreatitis. J Biol Chem 2014; 289:27551–27561.

46. Yang BS, Gilbert JD, et al. Overexpression of Myc suppresses CCAAT transcription factor/nuclear factor 1-dependent promoters in vivo. Mol Cell Biol 1993; 13:3093–3102.

47. Nebl G, Mermod N, et al. Post-transcriptional down-regulation of expression of transcription factor NF1 by Ha-ras oncogene. J. Biol Chem 1994; 269:7371–7378.

48. Denny SK, Yang D, et al. Nfib Promotes Metastasis through a Widespread Increase in Chromatin Accessibility. Cell 2016; 166:328–342.

49. Semenova EA, Kwon MC, et al. Transcription Factor NFIB Is a Driver of Small Cell Lung Cancer Progression in Mice and Marks Metastatic Disease in Patients. Cell Rep 2016; 16:631–643.

50. Lukas J, Pospech J. et al. Role of endoplasmic reticulum stress and protein misfolding in disorders of the liver and pancreas. Adv Med Sci 2019; 64:315–323.

51. Kawaguchi Y, Cooper B, et al. The role of the transcriptional regulator Ptf1a in converting intestinal to pancreatic progenitors. Nat Genet 2002; 32:128–134.

52. Rubio-Camarillo M, López-Fernández H, et al. RUbioSeq+: A multiplatform application that executes parallelized pipelines to analyse next-generation sequencing data. Comput Methods Programs Biomed. 2017; 138:73–81.

