## Supplementary Materials for "NFIC regulates ribosomal biology and ER stress in pancreatic acinar cells and suppresses PDAC initiation"

### SUPPLEMENTARY MATERIAL

### SUPPLEMENTARY FIGURES

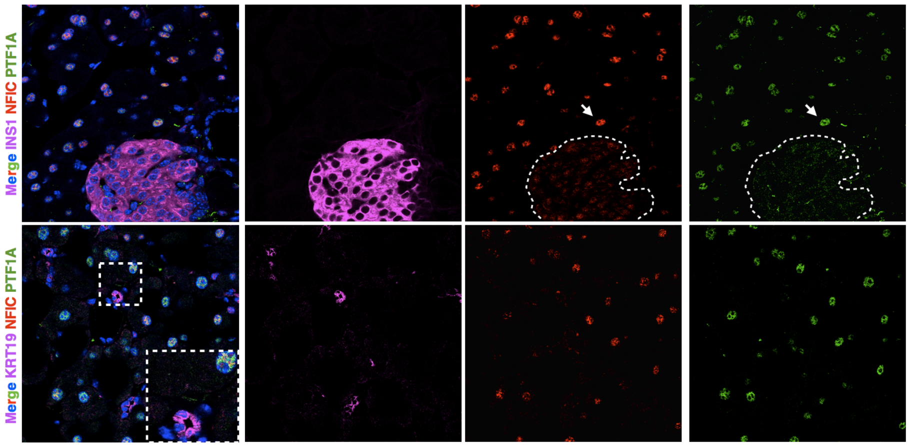

**Supplementary Figure 1. In the adult mouse pancreas, NFIC is expressed at highest levels in acinar cells and at lower levels in ductal and endocrine cells**. Expression analysis of NFIC in the pancreas of 8 week-old wild-type mice using triple IF on 4% PFA-fixed sections. NFIC is expressed at higher levels in acinar cells than in endocrine cells (INS1$^{+}$) and it is undetectable in ductal cells (KRT19$^{+}$). Square with dotted lines denotes the magnified area. Arrow denotes acinar cell. One representative image of 4 wild-type pancreata is shown.

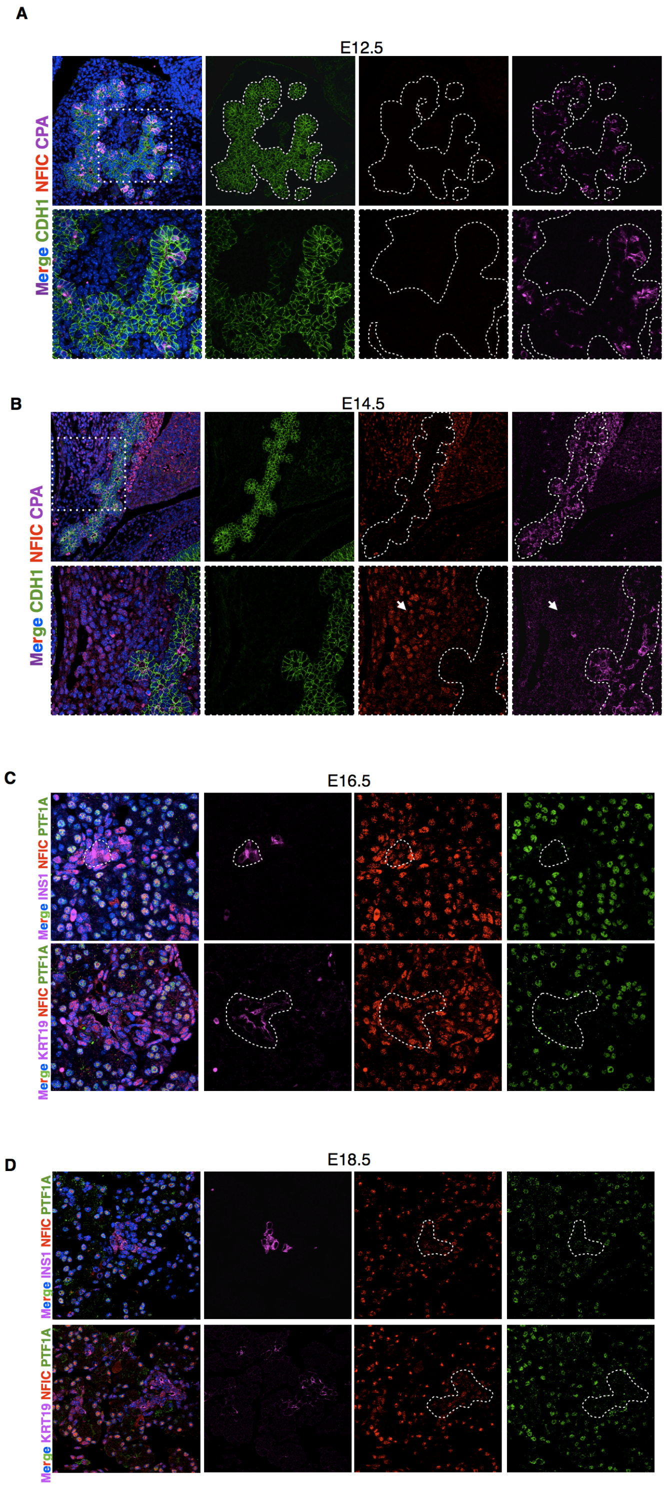

**Supplementary Figure 2. NFIC is expressed at late stages of pancreatic development**. (A,B) Expression analysis of NFIC in the pancreas of E12.5 (A) and E14.5 (B) wild-type embryos using triple IF on 4% PFA-fixed sections shows undetectable levels of NFIC. Expression of CDH1 and CPA was used to trace pancreatic progenitors. Broken lines delineate epithelial cells of the embryonic pancreas; arrows point to cells outside the embryonic pancreas showing the expression of NFIC in nonpancreatic progenitor cells in E14.5 embryos. (C,D) Expression of NFIC in the pancreas of E16.5 (C) and E18.5 (D) wild-type embryos using triple IF with antibodies detecting PTF1A, INS1, and KRT19. The expression of NFIC in acinar (PTF1A$^{+}$), endocrine (INS1$^{+}$) and ductal cells (KRT19$^{+}$) is shown. One representative image of 3 wild-type pancreata is shown. Square with broken lines denotes the region magnified.

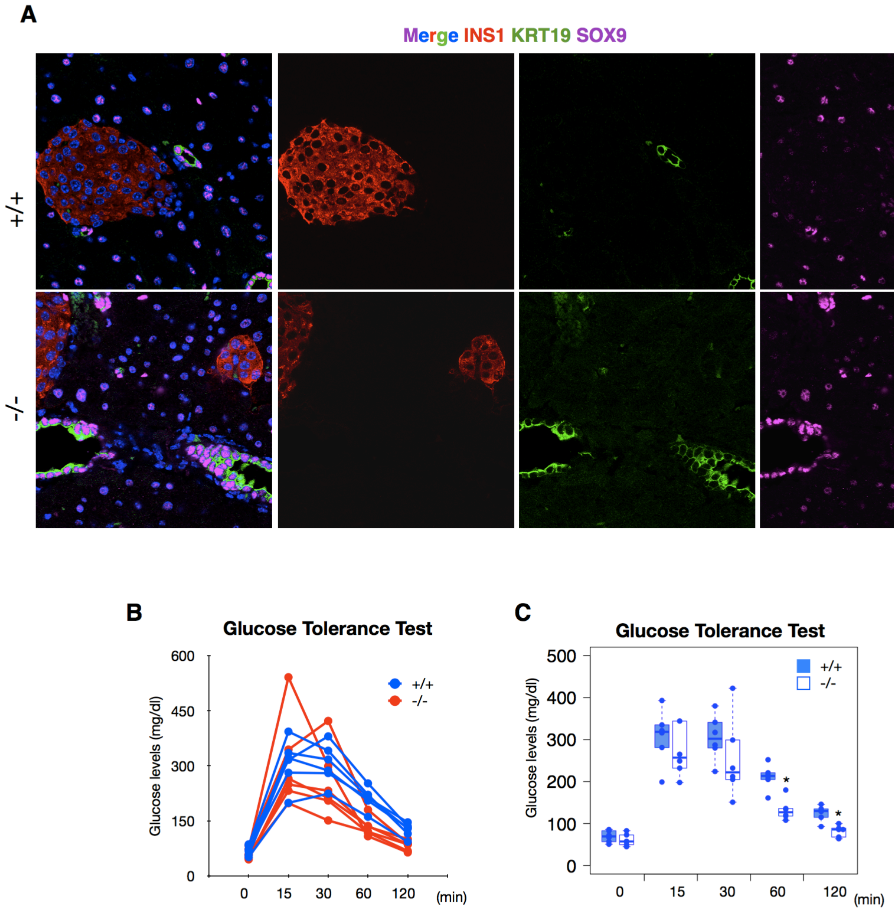

**Supplementary Figure 3. *Nfic$^{-/-}$* mice display no major abnormalities in ductal or endocrine cells and have a normal response to glucose overload**. (A) IF analysis of INS1, KRT19 and SOX9 expression in wild-type and *Nfic$^{-/-}$* pancreata (8-10 weeks). A representative image is shown. (B,C) Glucose tolerance test in wild-type and *Nfic$^{-/-}$* mice (11-17 weeks) showing similar glucose levels in mice of both genotypes at 0, 15 and 30 minutes but reduced glucose levels in *Nfic$^{-/-}$* mice at 60 and 120 min (n=6 male mice/genotype). Fasting glucose levels were measured before and after intra-peritoneal injection of glucose (2g/kg of body mass). Data for each individual mice (B) and grouped by genotype (C) are shown. In (C), two-sided Mann-Whitney U test was used to calculate statistical significance. P<0.05 (*). P<0.01(**).

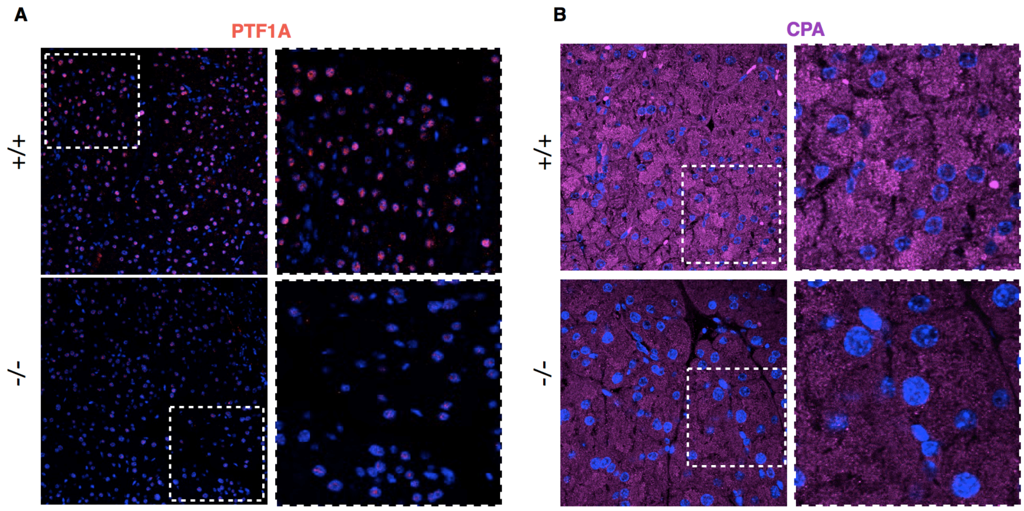

**Supplementary Figure 4. NFIC is required for normal acinar cell differentiation**. (A,B) IF analysis of PTF1A (A) and CPA (B) expression in wild-type and *Nfic*$^{-/-}$ pancreata showing regions of the exocrine parenchyma with reduced expression of PTF1A and CPA. One representative image is shown (n=3/group).

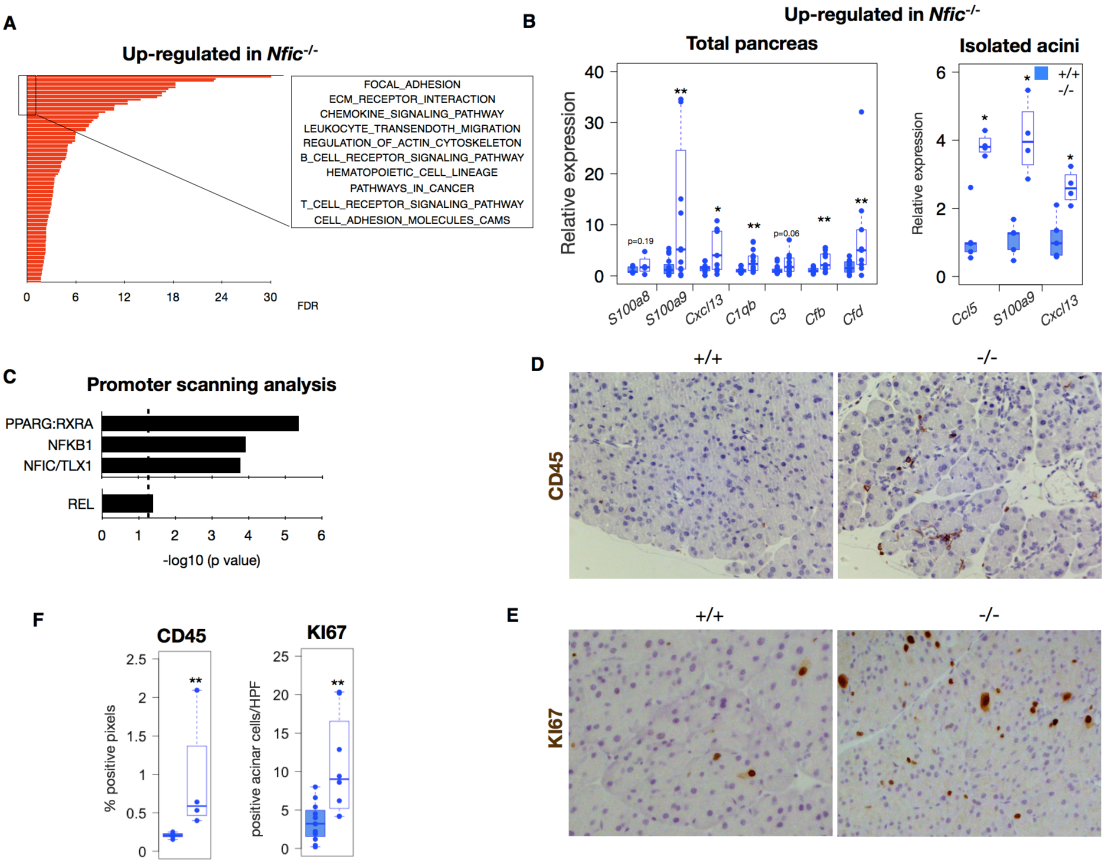

**Supplementary Figure 5. *Nfic*$^{-/-}$ pancreata show up-regulation of inflammatory gene transcripts, increased number of CD45$^{+}$ cells, and higher number of proliferating acinar cells than wild-type pancreas**. (A) Gene set enrichment analysis of the pre-ranked list of differentially expressed transcripts in the pancreas of *Nfic*$^{-/-}$ mice compared to wild-type. The pre-ranked list was computed with REACTOME and significant gene sets were ranked by the FDR value. Top-10 most significant are displayed. Up-regulated genes in *Nfic*$^{-/-}$ pancreata belong to inflammatory gene sets. (B) RT-qPCR of inflammatory transcripts in the pancreas (left panel) or freshly isolated acinar cells (right panel) from wild-type and *Nfic*$^{-/-}$ mice (n>4/group). One-sided Mann-Whitney U test was used to calculate statistical significance. P<0.05 (*). P<0.01(**). (C) Promoter Scanning analysis (PScan) of up-regulated genes in *Nfic*$^{-/-}$ pancreata showing the over-representation of putative PPARγ:RXRA, NFKB1, and NFIC motifs and the REL family of transcription factors, on the promoter of up-regulated genes in the *Nfic*$^{-/-}$ pancreata. Dotted line in C refers to threshold for statistical significance (-log10[0.05]=1.30). (D,E) Quantitative analysis of leukocyte infiltration (D) (n=3/group) and of proliferating acinar cells (E) in the pancreas of wild-type and *Nfic*$^{-/-}$ mice (n>5/group). For KI67 analysis, random images (n=10) were taken from each pancreas and positive acinar cells were considered. One-sided Mann-Whitney U test was used to calculate statistical significance. P<0.05 (*). P<0.01(**). (F) Quantification of data from panels D and E.

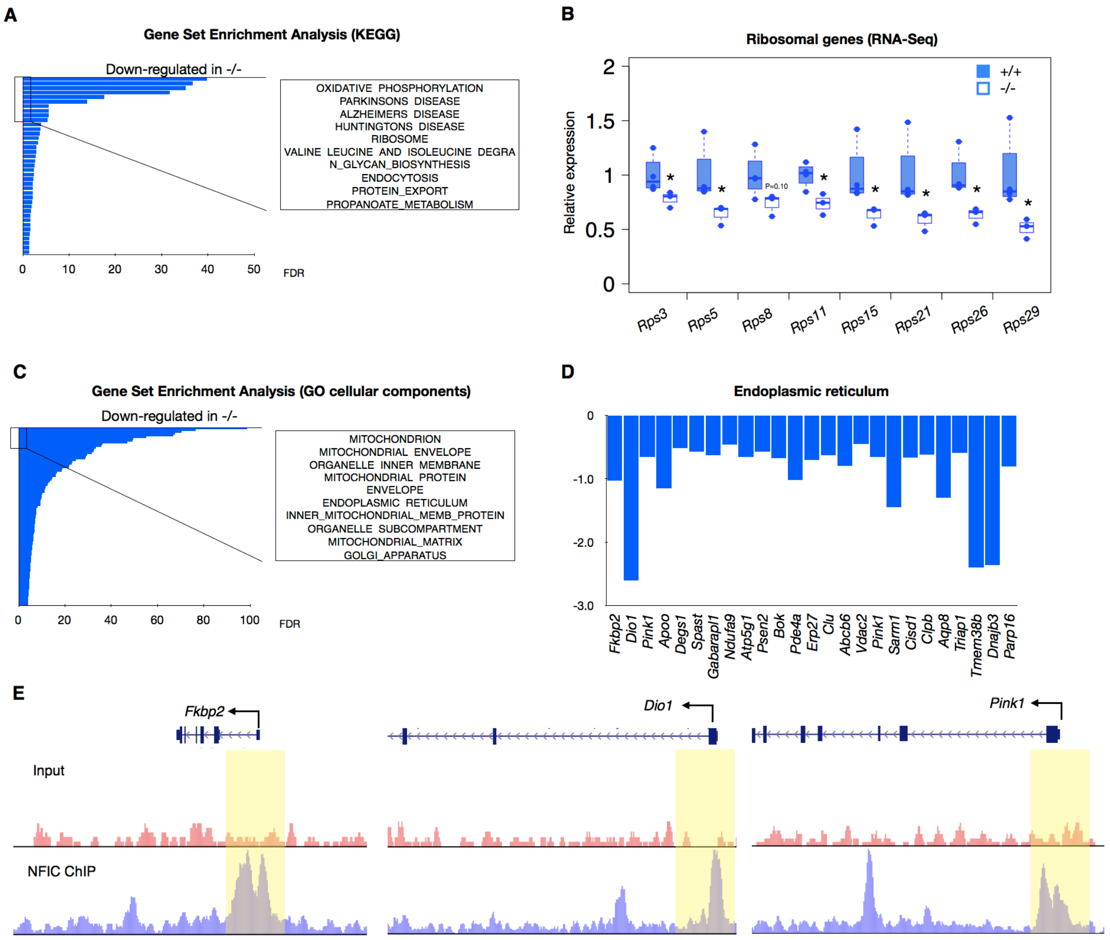

**F**

**NFIC ChIP-Seq (bp distance to TSS)**

| | GM12878 | ECC1 | HepG2 | SK-N-SH | K562 |
|---|---|---|---|---|---|
| *Rps5* | [-1461;+304] | [-760;+100] | [-836;+191] | - | - |
| *Rps11* | [-613;+379] | [-473;+379] | [-479;+255] | - | [-439;+4] |
| *Rps12* | [-1986;+375] | [-581;+187] | [-661;+295] | - | [-36;+124] |
| *Rps15* | [-328;+313] | [-313;+310] | [-398;+292] | - | [-76;+367] |
| *Rps16* | [-400;+470] | [-361;+230] | [-386;+224] | - | [-83;+360] |
| *Rps21* | [-300;+101] | [-612;+180] | [-498;+376] | - | - |
| *Rps26* | [-546;+588] | [-394;+590] | [-217;+461] | - | [+207;+317] |
| *Rps28* | [-367;+487] | [-378;+485] | [-723;+154] | - | [+10;+181] |
| *Rpl36* | [-680;+738] | [+836;+1419] | [-632;+79] | [-484;-20] | [-375;+68] |
| *Rpl37* | [-514;+590] | [-312;+313] | [-640;+465] | - | [-166;+135] |

**Supplementary Figure 6. NFIC is involved in mRNA and protein metabolism**. (A) GSEA of differentially down-regulated genes in *Nfic-/-* pancreata using KEGG. Significant gene sets were ranked by the FDR value and top-10 most significant gene sets are displayed. When multiple gene sets display a similar FDR, a representative gene set is shown (n>3/group). Down-regulated genes in *Nfic*$^{-/-}$ pancreata belong to mRNA, protein metabolism, oxidative phosphorylation and protein export. (B) Expression analysis by RNA-Seq of ribosomal gene transcripts showing down-regulation in *Nfic*$^{-/-}$ pancreata. Expression values were normalized to those in wild type pancreata (n>3/group). (C) GSEA of differentially down-regulated genes in *Nfic*$^{-/-}$ pancreata using GO cellular components showing enrichment in mitochondrion, endoplasmic reticulum and Golgi apparatus. (D) Expression analysis by RNA-Seq of genes coding for ER components showing down-regulation in *Nfic*$^{-/-}$ pancreata (FDR<0.05) (n$\geq$3/group). (E) Composites of NFIC ChIP-Seq showing a peak at the promoter of *Fkbp2*, *Dio1,* and *Pink1* as representative ER genes. (F) NFIC binding to ribosomal gene promoters in GM12878, ECC1, HepG2, SK-N-SH, and K562 cells. Data corresponds to distance to the transcription start site (TSS).

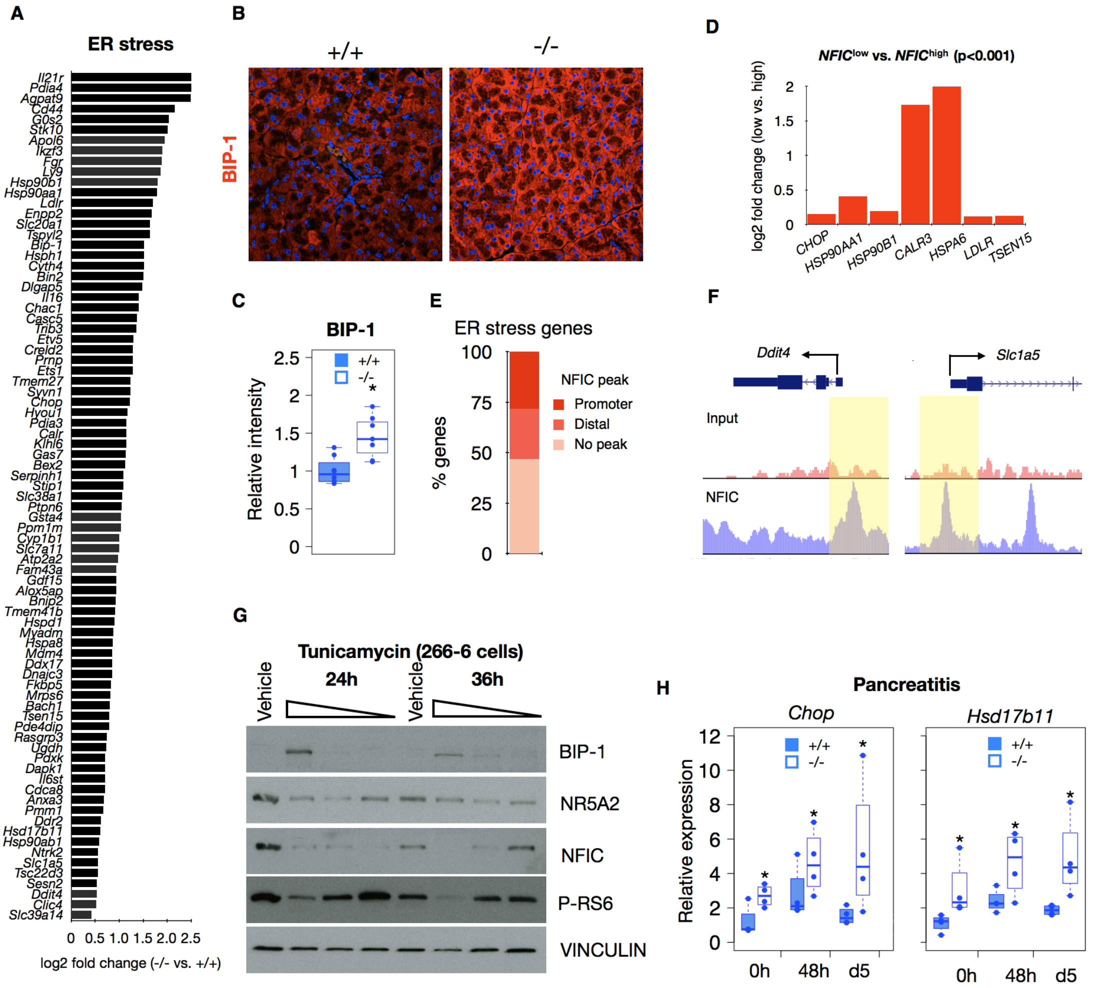

**Supplementary Figure 7. NFIC regulates aspects of the ER stress response**. (A) Expression of genes involved in UPR and ER stress showing an up-regulation in *Nfic*$^{-/-}$ pancreata (RNA-Seq). (B) IF analysis of BIP-1 in wild type and *Nfic*$^{-/-}$ pancreata (n>5/group). (C) Quantification of the BIP-1 expression intensity in wild type and *Nfic*$^{-/-}$ pancreata from panel B. Individual dots correspond to the average of at least 15 images for each pancreas. (D) Expression analysis showing up-regulation of ER stress genes in *NFIC*$^{low}$ compared to *NFIC*$^{high}$ human pancreata. (E) Bar graph showing the percentage of ER stress genes with an NFIC peak at the promoter or at distal regions. (F) Composites of NFIC ChIP-seq showing peaks at the promoter of *Ddit4* and *Slc1a5*. (G) 266-6 cells incubated with vehicle or increasing TM concentrations (10 nM, 1 nM, 0.1 nM) for 24h or 36h. Data shows the up-regulation of BIP-1, and down-regulation of NFIC and p-RS6, by 24/36h and the down-regulation of NR5A2 by 24h in 266-6 cells treated with TM (one representative experiment is shown). (H) RT-qPCR expression analysis showing up-regulation of *Ddit3* and *Hsd17b11* in *Nfic*$^{-/-}$ pancreata (n=4 mice/group). In all analyses, one-sided Mann-Whitney U test was used to calculate statistical significance. P<0.05 (*).

### SUPPLEMENTARY TABLES

**Supplementary Table 1. List of antibodies used.**

**Supplementary Table 2. List of primers used for RT-qPCR.**

**Supplementary Table 3. List of primers used for ChIP-qPCR.**

**Supplementary Table 1. List of antibodies used.**

| Protein | Catalog reference | Working concentration and (technique) |
|---|---|---|
| ACTIN | Sigma-Aldrich, MA1-744 | 0.02 μg/mL (WB) |
| BIP-1(HSPA5) | Cell Signalling, C50B12 | 0.8 μg/mL (IF); 0.2 μg/mL (WB) |
| Long ribosomal RNAs | Thermofisher MA1-13017 | 2 μg/mL (IF) |
| CD45 | Novus Biologicals, NB110-93609 | 0.8 μg/mL (IHC) |
| CDH1 | BD transduction, C20 820 | 0.25-0.35 μg/mL (IHC,IF) |
| CEL | Abcam, ab87431 | 0.2 μg/mL (WB) |
| CHOP (DDIT3) | Cell Signalling, CL63F7 | 0.2 μg/mL (WB) |
| CPA1 | RnD Systems, AF2765 | 1 μg/mL (IF) |
| CPA1 | Biorad (formerly AbD serotec), 1810-0006 | 0.5 μg/mL (WB) |
| CTRB1 | Biorad (formerly AbD serotec), 2100-0657 | 0.5 μg/mL (WB) |
| ERK | Cell Signalling, CST #9102 | 0.1 μg/mL (WB) |
| HA- tag | Sigma -Aldrich, F3165 | 0.1 μg/mL (WB) |
| Histone H3 | Abcam, ab1791 | 0.05 μg/mL (WB) |
| IgG (Goat) | Millipore, NI02 | |
| IgG (Mouse) | Santa Cruz, sc-2025 | |
| IgG (Rabbit) | Millipore, 12-370 | |
| INS1 | Dako, A0564 | 1/400 (IF) |
| KI67 | Leica, clone MM1, K2 | 0.05 μg/mL (IHC) |
| KI67 | Bethyl, IHC-00375 | 0.05 μg/mL (IHC) |
| KRT19 (Troma3 ) | Monoclonal Antibodies Unit (CNIO) | 1/25 (IF); 1/50 (IHC) |
| NFIC | Bethyl, A303-123A | 0.4 μg/mL (IHC on formalin-fixed sections) |
| NFIC | Abcam, ab89516 | 1.25 μg/mL (IHC/IF on PFA-fixed sections); 0.5 μg/mL (WB); 1 μg/ChIP or IP |
| NR5A2 | Everest, EB12283 | 2 μg/ChIP or IP; 0.5 μg/mL (WB) |
| P-EIF4E ($Ser^{209}$) | Cell Signalling, CST #9741 | 0.2 μg/mL (WB) |
| P-S6 ($Ser^{240/244}$) | Cell Signalling, CST #2215 | 1 μg/mL (IHC)(IF); 0.2 μg/mL (WB) |
| P-S6K1($Thr^{389}$) | Cell Signalling, CST #9205 | 0.2 μg/mL (WB). |
| PTF1A | Kindly provided by B. Bréant (INSERM). | 1/400 (IHC); 1/200 (IF); 1/1000 (WB); 1/500 (ChIP) |
| SOX9 | Millipore AB535 | 0.4 μg/mL (IF) |
| VINCULIN | Sigma -Aldrich, Clone hVIN-1 | 0.1-0.13 μg/mL (WB) |

**Supplementary Table 2. List of primers used for RT-qPCR.**

| Gene name (cDNA) | Forward | Reverse |
|---|---|---|
| *Amy2a5* | TGGCGTCAAATCAGGAACATG | AAAGTGGCTGACAAAGCCCAG |
| *Bip-1/Hspa5* | TCATCGGACGCACTTGGAA | CAACCACCTTGAATGGCAAGA |
| *Cel* | AAGTTGCCCGTGAAAAAGCAG | ATGGTAGCAAATAGGTGGCCG |
| *Cela1* | TGTGTCACACCCCTACTGGA | TTGTTAGCCAGGATGGTTCC |
| *Cela2a* | AGGTGGAGGATGATGTGAGC | TGTCAGAACCCAGTTGTTGG |
| *Cela3b* | AGTTGTCAATGGCGAGGAAG | CAGAACCCAGTCAGGGGTAA |
| *C1qb* | CACAGAACACCAGGATTCCA | CCCACTGTGTCTTCATCAGC |
| *Ccl5* | GCCTCACCATATGGCTCGGACA | CCTTGACGTGGGCACGAGGC |
| *Ccl5* | GTGCCCACGTCAAGGAGTAT | CCCACTTCTTCTCTGGGTTG |
| *Ccl7* | CCAACCAGATGGGCCCAATGCATCC | TCAGCGCAGACTTCCATGCCC |
| *Ccl9* | TGGGCCCAGATCACACATGCAAC | CGGCCTGGTACACCCACCAC |
| *Complement C3 (C3)* | AGTGCTACTGCTGCTGTTGG | GCCGTAGGACATTGGGAGTA |
| *Complement Factor B (CFB)* | CCGAGACCAAAAGATTGTCC | TCCCCATTTTCAAAGTCCTG |
| *Complement Factor D (CFD)* | TGCACAGCTCCGTGTACTTC | CTCCTGGCCACCCAGAAT |
| *Complement Factor P (CFP)* | TATGCCTTCCAGGAGCATGA | CCATAAGGACCATGCTGACC |
| *Chop/Ddit3* | GTCCCTAGCTTGGCTGACAGA | TGGAGAGCGAGGGCTTTG |
| *Cpa1* | TACACCCACAAAACGAATCGC | GCCACGGTAAGTTTCTGAGCA |
| *Ctrb1* | GCAAGACCAAATACAATGCCC | TGCGCAGATCATCACATCG |
| *Cxcl13* | GCCTCTCTCCAGGCCACGGTAT | AGCCATTCCCAGGGGGCGTA |
| *Hprt* | CGTCGTGATTAGCGATGATGA | ACAATGTGATGGCCTCCCA |
| *Hsd17b11* | CTGACTGCCTACGAATTTGCC | GTTTCCTCGATGCCGTTCTTATT |
| *Nfic* | TGGACCTGTACCTGGCCTAC | GCTCTCCTGGAAGTCTGTGG |
| *Nr5a2* | CGATCAGCGGGAGTTTGTAT | CATTCACCTGCTCTTGGACA |
| *Nr5a2* | GTTGAGTGGGCCAGGAGTAGTA | ACGCGACTTCTGTGTGTGAG |
| *Nr5a2* | TTGAGTGGGCCAGGAGTAGTA | ACGCGACTTCTGTGTGTGAG |
| *Nr5a2* | TCATGCTGCCCAAAGTGGAGA | TGGTTTTGGACAGTTCGCTT |
| *Pnlip* | ACAGATCAACACCCGCTTTC | CGGGTTTTTCTGTTTGTTCG |
| *Ptf1a* | AACCAGGCCCAGAAGGTTAT | AAAGAGAGTGCCCTGCAAGA |
| *Rnase1* | CAGCAGGACAAACAATGGAA | CCAATTCGTCTTGGAGTTCA |
| *Spink3* | GGCAACTAGCCTCTTTTCCA | GACAATGAAGGTGGCTGTCA |
| *Nr0b2* | AGCTGGGTCCCAAGGAGTAT | AGTGAGCCTCCTGTTGCAG |
| *Rps5* | CAAGCTCTTTGGGAAATGGA | GGGCAGGTACTTGGCATACT |
| *Rps8* | TTAGAAACCGGACCGTGAAG | TCTCAGGGCACGGTACTTCT |
| *Rps28* | CTCCTCTCCGCCAGATCG | GCCTTTGACATTTCGGATGA |
| *S100a10* | TCTTCGGCACTAGCCTCATC | ATTTGGGATGGCATTTTGAA |
| *S100a11* | CAAAAGTACAGCGGGAAGGA | CTTCTTCATCATGCGGTCAA |
| Spliced *Xbp1* | AAGAACACGCTTGGGAATGG | CTGCACCTGCTGCGGAC |

**Supplementary Table 3. List of primers used for ChIP-qPCR.**

| **Gene name (promoter)** | **Forward** | **Reverse** |
|---|---|---|
| *Cel* | CCATAAATACTGGAGAAGAGGAGAGAA | CAGGCGCCCCATGGT |
| *Cela2a* | GGCACTCTTCTCAGGGTGTT | GAGCATGTGTGTCCTTCCATT |
| *Cela2a* | GGACCTGTCTTTGGCATGTT | TTTCCATTCCCTTGTCGTTC |
| *Cpa1* | CATGGTCAAGGGTGAAAGC | GAGGCAGGAGTTCCAAACTG |
| *Cpa1* | AGCTGACCCCATGGTCAAGGG | GGGTCCCTGGGGACAGTTCCC |
| *Ctrb1* | CCAACCAGAAAGGTCCAAGA | CAGAGCAGCTGTCCTTTTCC |
| *Chop/Ddit3* | CCTCCCACCACCATCGAC | GAGGAGGTGAGTGAGTCATGC |
| *Hspa5/Bip-1* | TGTTTCTCCTTCACCCCAAG | GCGGAGAAAGGGAATAGGTT |
| *Hsp90aa1* | TACGCAACCAGAACCCCAGT | AGGCAAAAGGTCCCCCTTC |
| *Negative region* | TTGGGTGTTGGGAACTGAAT | CCCTTCTCTGCCTTCTGATG |
| *Nfic promoter #1* | AGGCTGCTAGCGCTGTTCTA | CACTGCCAAAGATGGAGGAG |
| *Nfic promoter #2* | CACCTGGCTGTCATGTGTC | CAATTGCATGTCAGGCTTGT |
| *Nfic promoter #3* | GGAGGAGGGGAGTGAGAGTT | ACACGACTCCTGCCTGTCTC |
| *Nr0b2* | GACAAGCTGACAGTCACACACTAGAA | GCCCTGGCACCTGGTTTA |
| *Pnlip* | CCGGTACTTCTCTGGGCTTA | CTACGGACAACGAGCAAACA |
| *Pnlip* | CGCTCAGCCTCCGGTACTT | GGCTCCAAGGACGTACTTCTAATT |
| *Rps5* | TTAAGGGTGCTGATTGTAACGA | AGTATTTTGAGACAAAGTCCGATT |
| *Rps8* | CTGTGAACAGAGCGTTGGAG | AGGCCAACCTGGTCTACAAA |
| *Rps28* | GGGAAGGATTTGGGTTCTGT | GGAGCTAAAGCCAGGCAATC |
